# Separating cognitive and motor processes in the behaving mouse

**DOI:** 10.1101/2023.08.23.554474

**Authors:** Munib A. Hasnain, Jaclyn E. Birnbaum, Juan Luis Ugarte Nunez, Emma K. Hartman, Chandramouli Chandrasekaran, Michael N. Economo

**Affiliations:** Department of Biomedical Engineering, Boston University, Boston, MA; Graduate Program for Neuroscience, Boston University, Boston, MA; Center for Neurophotonics, Boston University, Boston, MA; Department of Psychological and Brain Sciences, Boston University, Boston, MA; Department of Neurobiology & Anatomy, Boston University, Boston, MA; Center for Systems Neuroscience, Boston University, Boston, MA

**Author notes:** These authors contributed equally (Munib A. Hasnain, Jaclyn E. Birnbaum). Order determined via coin flip.

## Abstract

The cognitive processes supporting complex animal behavior are closely associated with ubiquitous movements responsible for our posture, facial expressions, ability to actively sample our sensory environments, and other critical processes. These movements are strongly related to neural activity across much of the brain and are often highly correlated with ongoing cognitive processes, making it challenging to dissociate the neural dynamics that support cognitive processes from those supporting related movements. In such cases, a critical issue is whether cognitive processes are separable from related movements, or if they are driven by common neural mechanisms. Here, we demonstrate how the separability of cognitive and motor processes can be assessed, and, when separable, how the neural dynamics associated with each component can be isolated. We establish a novel two-context behavioral task in mice that involves multiple cognitive processes and show that commonly observed dynamics taken to support cognitive processes are strongly contaminated by movements. When cognitive and motor components are isolated using a novel approach for subspace decomposition, we find that they exhibit distinct dynamical trajectories. Further, properly accounting for movement revealed that largely separate populations of cells encode cognitive and motor variables, in contrast to the ‘mixed selectivity’ often reported. Accurately isolating the dynamics associated with particular cognitive and motor processes will be essential for developing conceptual and computational models of neural circuit function and evaluating the function of the cell types of which neural circuits are composed.

## Introduction

Goal-directed behavior follows from an interplay between cognitive and motor processes in the mammalian brain. For example, while dribbling down the court, a basketball player must track the players around them, decide which play to run, plan their next move, execute fast and accurate movements, and flexibly adapt those movements according to the actions of other players. The neural representations of these cognitive and motor processes are often distributed across many of the same brain areas^1–6^ and engaged concurrently^7–9^. For these reasons, disentangling the neural signatures of specific cognitive and motor processes is challenging.

Many behavioral tasks are designed with this issue in mind, aiming to isolate constituent neural processes by separating them into discrete temporal epochs. For example, in sensory-guided decision-making tasks, subjects are often trained to withhold responses while stimuli are presented so that perceptual decision making is isolated in time from associated actions. Analogous paradigms have been developed to uncover the neural underpinnings of working memory^10–12^, motor planning^13,14^, contextual encoding^15,16^, reward prediction^17,18^, and other cognitive processes^19–22^.

These paradigms often rely on the assumption that experimental subjects withhold motor output during task epochs in which instructed responses are absent. However, uninstructed movements not required for task completion – such as changes in posture, facial expressions, and gaze – are commonly observed in rodents, humans, and non-human primates during learned behavioral tasks^23–29^. Importantly, uninstructed movements explain much of the variance observed in brain-wide neural activity^27,30–32^ and can be strongly correlated with task variables^12,28,33–38^.

These observations imply a serious challenge for experimental studies addressing a wide-ranging set of cognitive processes. If uninstructed movements are correlated with a latent cognitive variable of interest – a stimulus perceived, a decision made, a memory stored, or a motor plan formed – then the neural dynamics leading to, or resulting from, uninstructed movements can be easily misconstrued as responsible for that cognitive process. One common approach to isolate putative cognitive dynamics is to track and ‘regress out’ movements^24,30,39–44^, but this approach assumes that cognitive and motor processes are separable - that is, driven by independent neural dynamics. However, some cognitive processes may be inherently embodied such that their associated neural dynamics are linked to overt movement^45,46^. Regressing out neural dynamics associated with embodied movements would lead to the inadvertent removal of the precise dynamics one wishes to study. Whether cognitive and motor dynamics are separable remains an open question and likely depends on the brain area, behavior, and cognitive process of interest. Effective methods both for evaluating separability within specific experimental paradigms and for effectively dissociating cognitive and motor dynamics, when separable, are lacking.

Here, we address the separability of dynamics associated with cognitive processes and correlated movements. We developed a behavioral paradigm in which mice perform sensory-guided movements involving multiple cognitive signals in which they often exhibit idiosyncratic, task-correlated uninstructed movements. We build upon work demonstrating how different neural processes can be multiplexed in a single brain region^39,40,47–50^ to develop a novel method for assessing whether cognitive dynamics can be separated from those associated with movements – and for isolating each component, when they are separable. This approach is simple to adopt and does not require tracking or segmentation of body parts. It also does not require an explicit choice of models relating neural dynamics to movement, avoiding common assumptions of linearity. We find that some cognitive signals are separable from dynamics associated with co-occurring movements while others are largely inseparable. When dynamics are separable, examining the component of neural dynamics unrelated to movement revealed trajectories that differed in notable ways from estimates of the same dynamics when corrupted by uninstructed movements. Strikingly, we found that cognitive and motor dynamics were largely encoded by separate populations of cells when uninstructed movements were accounted for. Together, these results highlight the importance of critical consideration of the relationship between cognition and movement to better understand the neural dynamics supporting complex processes and how they map onto the myriad cell types comprising neural circuits.

## Results

### Task-switching behavioral paradigm

We first designed a behavioral paradigm associated with multiple cognitive processes. In this paradigm, head-fixed mice performed two directional licking tasks that alternated block-wise within each behavioral session. These tasks varied in their cognitive demands but required the same instructed motor output.

This paradigm employed an established delayed-response task (DR) in which motor planning is temporally dissociated from movement execution^14^ (**Fig. 1a**). In the DR task, an auditory stimulus indicated the location of a reward. After a brief delay epoch with no auditory stimuli, a separate auditory ‘go cue’ instructed the mouse to move. If the mouse performed a directional tongue movement to the correct target, a water reward was delivered. In separate blocks of trials, mice performed a water-cued (WC) task in which all auditory stimuli were omitted. Instead, a drop of water was presented at a random point in time at a randomly selected reward port. Animals consumed water upon its presentation. Mice received no explicit cues signaling DR and WC blocks. The DR task required a perceptual decision that guides the planning of a subsequent motor action. In the WC task, uncertainty in the timing and location of reward prevented formation of a motor plan prior to reward presentation, and instead, mice detected reward availability using olfactory and/or vibrissal cues^51^. This structure required animals to maintain an internal representation of the current block identity (context) to maintain high task performance.

**Figure 1.**
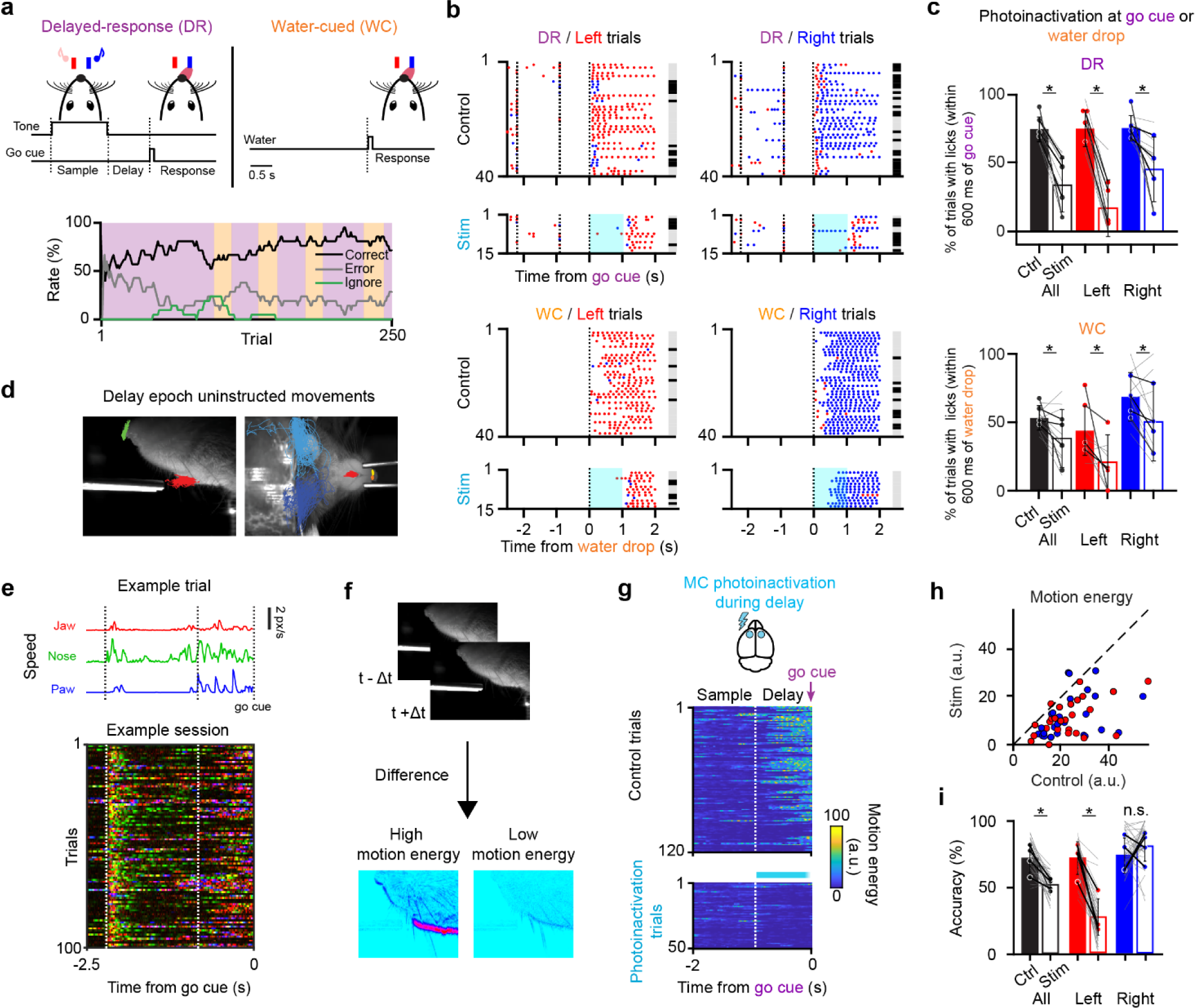
Cortical dependence and uninstructed movements during a two-context task. **a.** *Top*, Schematic of the two-context task. In delayed-response (DR) trials, an auditory stimulus was presented during the sample epoch (1.3 s) instructing mice to lick for reward to the right (white noise) or left (8 kHz tone). Mice were trained to withhold their response during a delay epoch (0.9 s) and initiated their response following an auditory go cue. In WC trials, mice were presented with a water reward at a random port at random inter-reward intervals. The WC task was introduced once mice became experts in the DR task. *Bottom*, 20-trial trailing average of correct, error, and ignore rates for an example session. **b.** Performance for an example session separated by trial type (left vs. right), control vs. photoinactivation trials, and by context. Photoinactivation (1 s) was initiated at the go cue. Blue dots indicate right licks and red dots indicate left licks. Grey bars, correct trials; black bars, error trials. **c.** Percent of trials with correct licks within 600 ms of the go cue on photoinactivation trials. Colored points represent mean values across animals (*n* = 4, individual animals connected by dark lines). Light gray lines denote individual sessions (*n* = 10). Bars are across-session means. Asterisks denote significant differences (*p* < 0.05) between control and photoinactivation trials (All DR trials: *p* = 0.0001; DR left trials: *p* = 4.5e-06; DR right trials: *p* = 0.0074; All WC trials: *p* = 0.0371; WC left trials: *p* = 0.0134; WC right trials: *p* = 0.0316, paired *t-*test). Error bars denote standard deviation across sessions. **d.** Behavior was tracked with high-speed video from side (*left*) and bottom (*right*) views. Trajectories of delay epoch uninstructed movements from an example session overlayed. **e.** Uninstructed movements are highly variable across trials and across time. *Top*, jaw, nose, and paw speed for an example trial. *Bottom*, feature overlay for a subset of trials in an example session. At each time bin, *t*, an [r, g, b] color value was encoded as [jaw*_t_*, nose*_t_*, paw*_t_*]. **f.** Schematic of motion energy calculation. Example frames depicting high and low motion energy are shown. **g-i.** Bilateral motor cortex photoinactivation during the delay epoch. **g.** Motion energy on control (*top*) and photoinactivation (*bottom*) trials for an example session. On photoinactivation trials, light was delivered for 0.8 seconds beginning at the start of the delay epoch. **h.** Average delay epoch motion energy on single sessions for control and photoinactivation trials (*n* = 29 sessions, 4 mice). Red points indicate left trials and blue points indicate right trials for a single session. **i.** Performance for control and delay epoch photoinactivation trials. Mice often defaulted to a right choice following delay epoch photoinactivation. Asterisks denote significant differences (*p* < 0.05) between control and photoinactivation trials (n=29 sessions, 4 mice). Black lines and points indicate averages across sessions for individual animals and light grey lines indicate sessions.

We examined uninstructed movements in this behavioral paradigm using high-speed video. We tracked movement of the tongue, jaw, nose, and paws^52^ and calculated instantaneous motion energy to quantify movement in a feature-agnostic manner^31^ (**Fig. 1d-f**). We found that mice performed uninstructed movements that varied in their identity and timing across both trials and contexts (**Fig. 1e** and **Supplementary Movie 1**).

Prior work has established the necessity of the antero-lateral motor cortex (ALM) and tongue-jaw motor cortex (tjM1) for the planning and execution of tongue movements^14,53–56^. To assess cortical involvement in the initiation of instructed movements (directional licking) in both behavioral tasks, we used optogenetic photoinactivation to inhibit the ALM and tjM1 at the go cue (DR) or water presentation (WC) bilaterally in VGAT-ChR2-EYFP mice (10 sessions, 4 mice). Simultaneous photoinactivation of both regions at the go cue (DR task) or water drop (WC task) impaired the initiation of movement in both contexts (**Fig. 1b,c**; DR trials: 41 ± 20% reduction, mean ± s.d., *p* = 1x10^-4^; DR left trials: 58 ± 19%, *p* = 4x10^-6^, DR right trials: 30 ± 28%, *p* = 7x10^-3^; All WC trials: 15 ± 20%, *p* = 0.04; WC left trials: 23 ± 24%, *p* = 0.01, WC right trials: 18 ± 23%, *p* = 0.03; paired *t*-tests; **EDFig. 1c,d**). To assess cortical involvement in the expression of uninstructed movements, and whether these movements might be controlled by ALM or tjM1 preferentially, we bilaterally silenced the ALM and tjM1 during the delay epoch of DR trials individually (ALM: 15 sessions, tjM1: 9 sessions, 4 mice). Uninstructed movements were suppressed by photoinactivation of either area (**EDFig. 1b**; ALM: ; 53 ± 31% reduction, mean ± s.d., *p* = 3x10^-4^; tjM1: 62 ± 24% reduction, *p* = 2x10^-3^, paired *t*-test) and more strongly suppressed by concurrent photoinactivation of both areas (**Fig. 1g-h**; 29 sessions, 4 mice; 73 ± 35% reduction, mean ± s.d., *p* = 7x10^-9^, paired *t*-test), similar to previous observations^34^ (but see ref^57^). In the DR task, delay photoinactivation of both areas led to a significant decrease in overall performance (**Fig. 1i**; 22% ± 14% reduction, mean ± s.d., *p* = 2x10^-6^ paired *t*-test; **EDFig. 1b**) as expected from the disruption of choice/planning dynamics^14^, often leading to animals responding consistently to a preferred side rather than in a manner commensurate with the sensory cue. These results indicate that both the ALM and tjM1 are involved in the generation of uninstructed movements and the planning and execution of instructed directional tongue movements in the two-context paradigm.

### Neural dynamics related to task variables

While movement- and task-related information is encoded in both the ALM and tjM1 in the DR task^53,58^ and other directional licking tasks^55^, task variables are more strongly represented in the ALM^55,56,59^ and so we focused our subsequent analyses there. We recorded activity extracellularly in the ALM with high-density silicon probes during the two-context paradigm to track dynamics associated with planning, context, and the execution of movements (**Fig. 2**; two-context paradigm: 12 sessions, 6 mice, 522 units including 214 well-isolated single units; an additional 3 mice were trained only on the DR task totaling 25 sessions, 9 mice, 1651 units including 483 well-isolated single units included in analyses of the DR task only; see **EDFig. 2a** for per-session statistics). In the DR task, we found that choice selectivity (right vs. left DR trials) was widespread across all task epochs as expected (sample: 36%; delay: 42%; response: 58% of 483 single units, 25 sessions, 9 mice, 483 single units; see **Methods**). Individual units showed a variety of activity patterns across trial types, including delay epoch preparatory activity preceding instructed movements, often taken to be a signature of motor planning^54,60^ (**Fig. 2b left**). Individual units were also selective for behavioral context during all task epochs, including during the inter-trial interval (ITI) (39% of single units, 12 sessions, 6 mice, 214 single units; see **Methods**), suggesting persistent coding of context (**Fig. 2b**, *right*).

**Figure 2.**
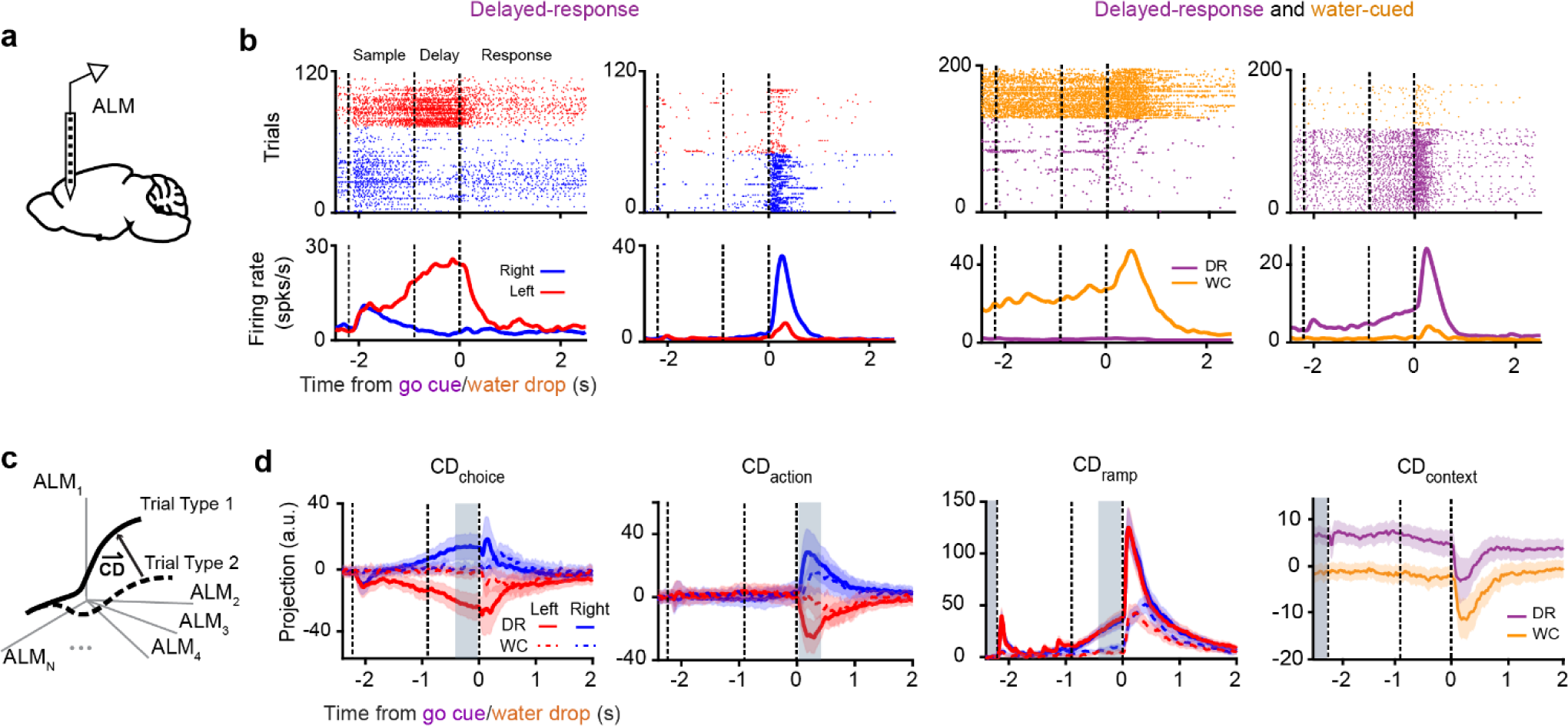
Single cell activity and population dynamics encode task-relevant cognitive and motor processes. **a.** Schematic of silicon probe recordings in the anterolateral motor cortex (ALM). **b.** Spike rasters (*top*) and peri-stimulus time histograms (PSTH; *bottom*) for four example units in the ALM. Units show selectivity for trial type (left vs. right) and context (DR vs. WC) during all task epochs. PSTHs for DR and WC tasks shown on right are averages of equal numbers of left and right trials. **c.** Schematic illustrating identification of coding directions (CD). Coding directions are defined as directions in state space that maximally separate activity between trajectories defined by trial types (CD_choice_, CD_action_), context (CD_context_), or time points (CD_ramp_). **d.** Projections along each of four defined CDs. Gray shaded regions indicate time points used for CD estimation. Mean and 95% confidence intervals of bootstrapped distributions shown.

We next sought to understand how the selectivity patterns observed in individual cells were encoded at the population level. We examined neural activity in state space, in which the firing rate of each unit represents one dimension. We defined one-dimensional coding directions (CD) within the high-dimensional state space that best encoded task variables^54,61–63^ (**Fig. 2c**). We first examined putative neural correlates of three cognitive processes – choice, urgency, and contextual encoding.

We defined a coding direction for left-right trial type, CD_choice_, as the direction in state space that best differentiates activity on left versus right trials during the delay epoch of DR trials. Projections along CD_choice_ revealed that choice selectivity in the ALM emerges early in the sample epoch and continues to grow throughout the delay epoch before decaying at the onset of the response epoch (**Fig. 2d**). Choice could be reliably decoded from projections along CD_choice_ across sessions (**EDFig. 2c**; AUC: 0.86 ± 0.11 mean ± s.d.). Such preparatory activity is predictive of animals’ upcoming instructed responses^13,54,58,61,64,65^ and has been taken to represent a neural correlate of a motor plan^54,60^ or choice^32,62,66^. We next defined CD_ramp_ as the direction that captures trial-type independent increases or decreases in preparatory activity between the onset of the stimulus and the go cue in DR trials^61,62,67,68^. This non-selective ramping has been thought to represent an urgency or timing signal^69–72^. Finally, we defined CD_context_, the direction that best differentiates activity during the inter-trial interval of DR and WC trials (see **Methods**). Projections along CD_context_ revealed that context is encoded in the ALM throughout each trial (**Fig. 2d**) with strong modulation at the onset of the response epoch.

In addition to neural activity putatively related to these cognitive processes, we also examined neural signals associated with the execution of movements. We defined an additional coding direction, CD_action_, as the direction that captures trial-type selective changes in neural activity that emerge after the go cue. Projections of neural activity along CD_action_ robustly encoded movement direction in the ALM, as expected. Together, these four coding directions explained 53 ± 11% (mean ± s.d.) of the variance in trial-averaged neural data (**EDFig. 2b**) We focus our analyses on these four population-level signals, which putatively encode ALM dynamics associated with cognitive and motor processes in the two-context task-switching paradigm.

### Uninstructed movements related to preparatory dynamics

We next examined uninstructed movements in the DR task and their correlation with both task variables and preparatory dynamics (**Fig. 3**). We found that animals frequently performed uninstructed movements that were correlated with trial type (**Fig. 3a**) and time within each trial. On single trials, upcoming choice could be decoded from uninstructed movements beginning immediately after the presentation of the sample tone, with accuracy increasing throughout the delay epoch (**Fig. 3b**).

**Figure 3.**
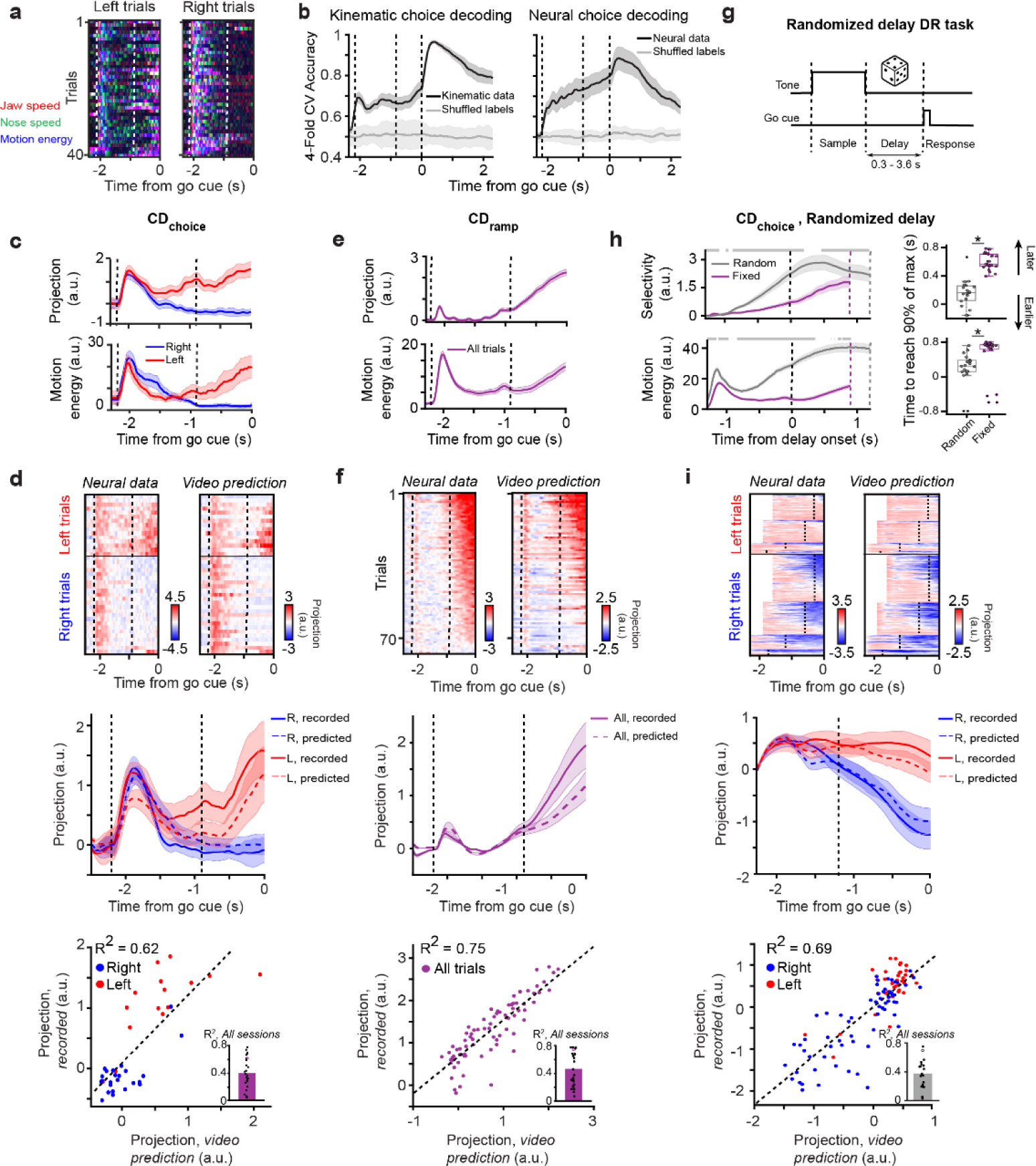
Uninstructed movements are tightly linked to putative planning dynamics. **a.** Choice-dependent uninstructed movements. Overlayed trajectories from a random subset of DR trials in an example session. In this example, uninstructed movements are prominent on left, but not right, trials during the delay epoch. **b.** Choice (left vs. right) decoding from kinematic features and neural population data. Gray lines denote shuffled choice labels. **c.** *Top*, average projection onto CD_choice_ on left (*red*) and right (*blue*) trials for the example session depicted in (a). *Bottom*, average motion energy for all correct left and right trials. **d.** Observed and video predictions of single-trial projections of neural activity onto CD_choice_ (see **Methods**) for the same example session as in (a). *Top*, Heatmap of observed and predicted single-trial projections. Trials are sorted by the observed average projection magnitude in the late delay, with left and right trials sorted separately. *Middle*, trial-averaged CD_choice_ projections and predictions. *Bottom*, scatter plot of the average delay epoch projection of neural data onto CD_choice_ versus corresponding video predictions. Dots denote single trials and dashed line denotes linear fit. *Inset*, R² values for all sessions (*n* = 25 sessions). Open circle denotes the example session in (a). **e.** Same as (c) but for CD_ramp_ (different example session). Left and right hits are grouped (*purple*). **f.** Same as (d) but for CD_ramp_. Same session as (e). **g.** Schematic of the randomized delay task. The delay epoch duration was randomly selected from six values (see **Methods**) with frequencies following an exponential distribution. **h.** Differences in choice selectivity and uninstructed movements between the randomized and fixed DR tasks. *Top left*, selectivity in CD_choice_ projections averaged across sessions for randomized delay (*gray*; *n* = 19 sessions) and fixed delay (*purple*; *n* = 25 sessions) tasks. Vertical lines indicate time of delay onset (*black*), go cue for the fixed delay (*purple*) and or go cue for the randomized delay (*gray*). Only trials with a delay duration of 1.2 s are shown for the randomized delay task for clarity. Gray bar at the top denotes timepoints where the slopes of the curves are significantly different (*p* < 0.05, two-sided *t*-test). *Bottom left*, same but for motion energy. *Top right*, time relative to delay onset that selectivity in CD_choice_ projections reaches 90% of its maximum value. Colored dots, individual sessions; black dots, outlier sessions. Asterisk denotes significant difference (*p* < 0.05, two-sided *t*-test) between latencies in randomized vs. fixed delay sessions. *Bottom right*, same but for session-averaged motion energy. **i.** Same as (d) for an example session of the randomized delay task. *Top*, trials with delay durations of 0.3, 0.6, 1.2, and 1.8 s are shown. *Middle* and *bottom*, trials with delay duration of 1.2 s only. Lines with shaded regions depict mean and 95% confidence intervals of bootstrapped distributions throughout.

Not only were uninstructed movements related to the animal’s upcoming actions, but they were highly correlated with preparatory dynamics on a trial-by-trial and moment-by-moment basis. The onset and magnitude of uninstructed movements often coincided with the onset and magnitude of choice selectivity and ramping (**Fig. 3c,e**). We used cross-validated multiple linear regression to predict projections along CD_choice_ and CD_ramp_ on a trial-by-trial basis from captured kinematic features of movement. Model predictions were often highly correlated with motor planning signals, indicating a close relationship between animals’ uninstructed movements and preparatory dynamics (**Fig. 3d,f**; CD_choice_ : R^2^ = 0.41 ± 0.23, mean ± s.d across sessions, CD_ramp_ : R^2^ = 0.48 ± 0.23, *n* = 25 sessions), although this relationship was variable across animals and sessions (**Fig. 3d,f,i bottom row**) and variable in the kinematic features that were most predictive (**EDFig. 3a-c**). This variability is consistent with other studies^73^ and highlights the need for analytical methods for assessing the relationship between neural signals and related movements on a session-by-session basis.

To further understand the relationship between uninstructed movements and preparatory dynamics, we trained additional animals on a randomized delay task in which the timing of the go cue cannot be anticipated^65^. Incorporating uncertainty into the length of the delay epoch leads to a qualitative change in population-level choice selectivity. Selectivity in CD_choice_ projections emerges earlier than in the standard fixed-delay DR task (**Fig. 3h** *top right*; fixed: 0.61 ± 0.11 s from delay onset, *n* = 25 sessions; randomized: 0.16 ± 0.19 s, *p* = 2.4x10^-12^, *n* = 19 sessions, two-sided *t*-test; **Fig. 3h**, *top left*), suggesting that motor plans are prepared prior to the earliest possible go cue time^65^. We found that the timing of uninstructed movements, too, shifted in an analogous fashion (**Fig. 3h**, *bottom*; fixed: 0.58 ± 0.43 s from delay onset; randomized: 0.19 ± 0.39 s, *p* = 0.01, two-sided *t*-test). Projections along CD_choice_ remained predictable from uninstructed movements on a single-trial level (**Fig. 3h**; average R^2^ across sessions: 0.38 ± 0.20, *n* = 19 sessions), providing further evidence of the tight link between preparatory dynamics and uninstructed movements.

### Uninstructed movements related to behavioral context

We next compared uninstructed movements across task blocks in the two-context paradigm (**Fig. 4**). We found that animals perform qualitatively different uninstructed movements in each behavioral context, despite both contexts requiring instructed movements to the same targets (**Fig. 4a**). Context could be decoded from kinematic features of movement across epochs – including during the ITI, in which external contextual cues were absent (**Fig. 4b**, *left*). The time-course of context decoding from neural population data and from movement kinematics was also similar (**Fig. 4b**, *right*).

**Figure 4.**
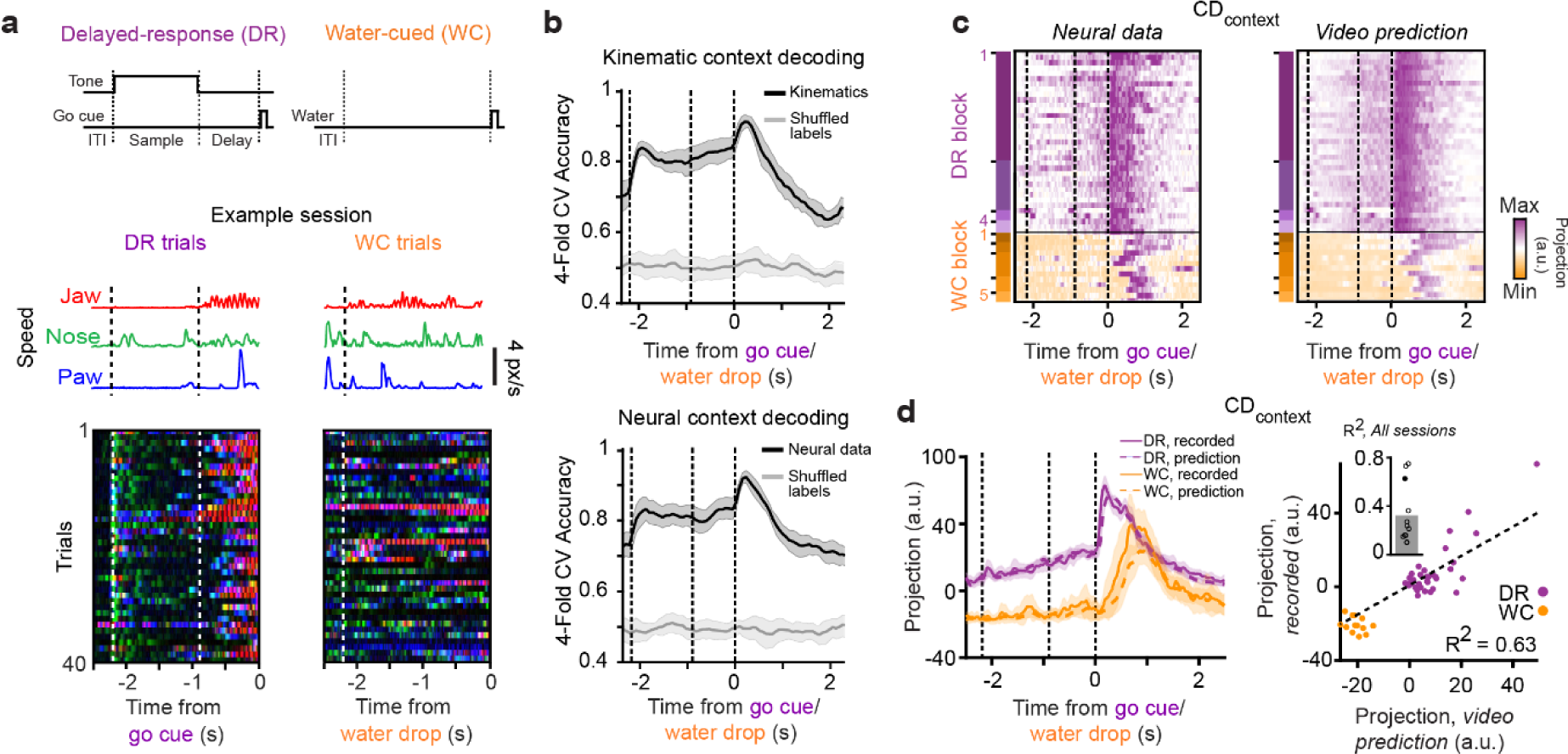
Uninstructed movements are closely related to the neural encoding of context. **a.** Schematic of tasks (*top*) and context-dependent uninstructed movements. Jaw/nose/paw movements on an example trial (*middle*) and overlayed trajectories of uninstructed movements across an example session (*bottom*) for DR task (*left*) and WC task (*right*). **b.** Context decoding from high-speed video (*top*) and neural data (*bottom*) as a function of time across all sessions. Gray lines denote shuffled context labels. **c.** *Left*, heatmap of single-trial projections of neural data onto CD_context_. *Right*, CD_context_ projections predicted from video in an example session. The chronological DR or WC block within the session is denoted by purple and orange rectangles, respectively, on the left of each plot. **d.** *Left*, trial-averaged projection of neural data onto CD_context_ and predictions from video (*dotted lines*) for the example session illustrated in (c). *Right*, scatter plot of average projection of neural data onto CD_context_ during the ITI versus average video prediction. Dots denote single trials; dashed line, line of best fit. *Inset*, R² values for all sessions (*n* = 12 sessions). Filled circle denotes the example session in (c). Lines with shaded regions depict mean and 95% confidence intervals of bootstrapped distributions throughout.

We sought to understand whether trial-to-trial variability in the neural representation of context-encoding could be predicted from uninstructed movements. We again trained a multiple linear regression model to predict single-trial projections along CD_context_ from kinematic features and found that, like putative choice and urgency dynamics, context-selective signals could be predicted with high fidelity (**Fig. 4c,d**; R^2^ = 0.49 ± 0.22, *n* = 12 sessions).

Taken together, these findings suggest that dynamics along CD_choice_, CD_ramp_, and CD_context_ could be associated with uninstructed movements and/or the cognitive processes related to the anticipation and planning of upcoming instructed movements and encoding of task context.

### Subspace decomposition of neural activity

We next examined whether neural signals encoding task variables could be isolated from movement-related signals, despite their correlation in time. We hypothesized that if the neural dynamics associated with cognitive and motor processes are separable, their components in the ALM should be driven by distinct latent signals.

Previous theoretical work demonstrated a simple mechanism that explains how neural activity can vary dynamically while ensuring that specific variables remain stably encoded^50^. In this formalism, a variable of interest may be decoded from a neural population, perhaps by a downstream circuit, through a linear transformation matrix, ***W***, which could be taken to follow from a particular pattern of synaptic connectivity. If the dimensionality of neural activity is larger than the dimensionality of the output signals decoded (i.e. if ***W*** is not square), then ***W*** will have a null space - dimensions along which neural activity can vary without affecting the decoded output^50^.

Applying this idea to recordings from the primate motor cortex demonstrated that motor planning dynamics are confined to the null space of a linear transformation between motor cortical activity and muscle activation^39^, thereby allowing motor planning signals to evolve dynamically in a manner that is independent of motor output. The orthogonal complement to this output-null subspace is the output-potent subspace; changes in activity along output-potent dimensions were then associated with changes in muscle activation^39^.

Subsequent work identified analogous subspaces of neural activity in the primate motor cortex as the orthogonal subspaces that capture the most variance in neural activity during the delay and response epochs of a delayed-response task to capture preparatory and movement-related activity, respectively^40^. This approach is effective when preparatory and movement-related dynamics are confined to distinct temporal epochs and when neural activity is exclusively related to motor preparation and motor execution, respectively, in those epochs. This approach is convenient in that it avoids the need to measure muscle activation or any other descriptor of motor output. Importantly, it also avoids the need to explicitly model the transformation between neural activity and motor output as a linear transformation. This transformation is, in general, unknown and can be highly nonlinear, particularly for movements directly controlled by central pattern generators, such as those that support much of the behavioral repertoire of rodents (e.g. locomoting, breathing, whisking, licking, chewing, vocalizing, swallowing, etc.).

We sought to determine if this computational framework could be used to isolate the neural dynamics associated with instructed and uninstructed movements from dynamics related to the encoding of choice, urgency, and context in our paradigm. Following the approach of Elsayed et al.^40^ we identified subspaces that maximally capture variance in trial-averaged neural activity in the delay and response epochs of our DR trials (**Fig. 5a** and **Methods**). We found that dynamics within ‘delay’ and ‘response’ subspaces were confined to the delay and response epochs, respectively, closely resembling results observed in primates^40^ (**Fig. 5b-d**). This outcome is expected when trial-averaged dynamics in different epochs occupy subspaces that are largely orthogonal^40^. However, activity within both subspaces remained highly correlated with uninstructed movements on single trials (**Fig. 5b,e,f**), suggesting that movement-related dynamics were not restricted to the ‘response’ subspace using this analytical framework. This follows from the assumption that planning- and movement-related processes are strictly confined to distinct task epochs, which was not the case in our paradigm. Uninstructed movements were expressed across all task epochs (**Figs. 1,3**, and **4**).

**Figure 5.**
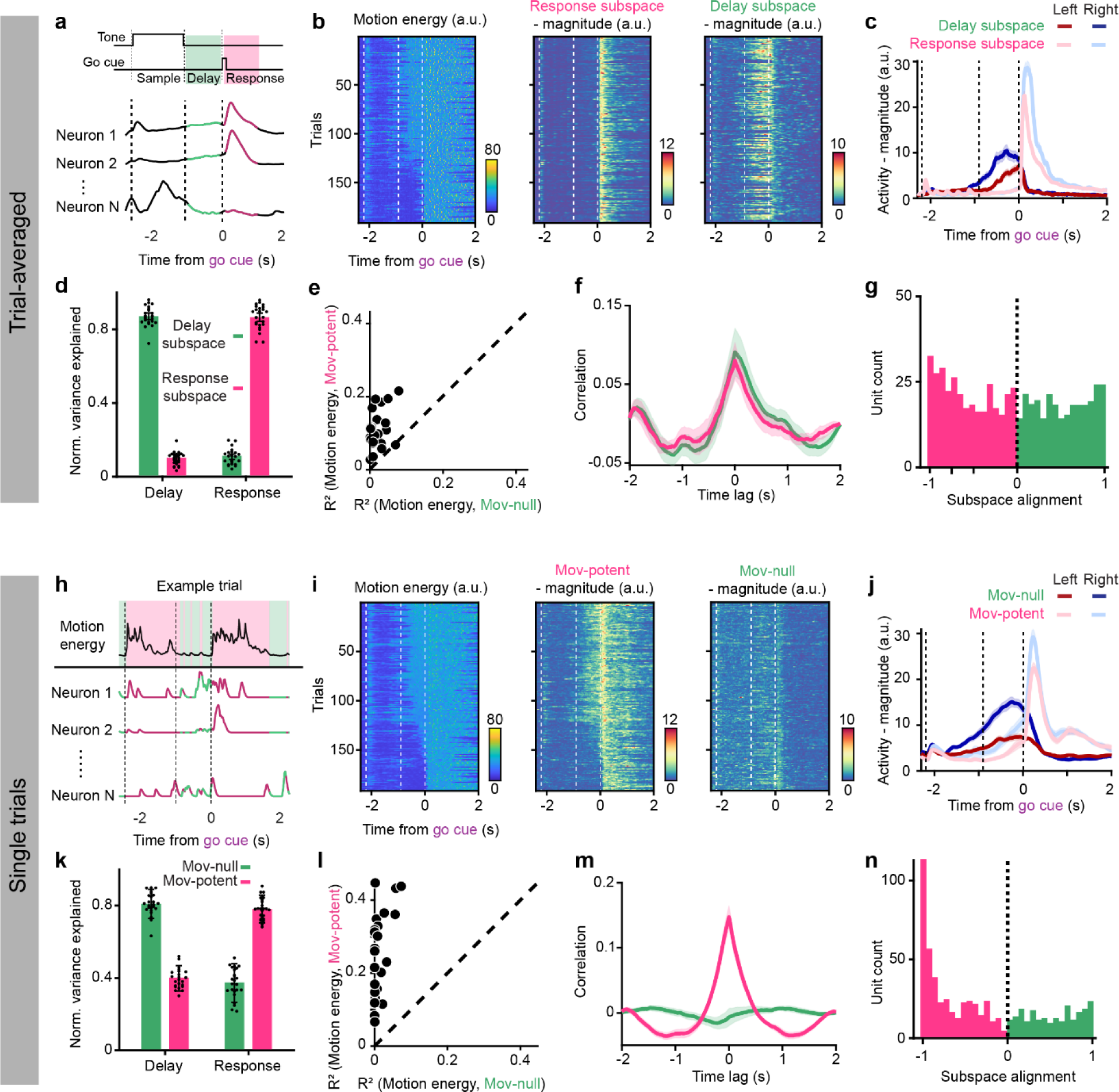
Subspace decomposition of neural activity using trial-averaged and single-trial data. **a.** Schematic of the approach for estimating delay (*green*) and response (*pink*) subspaces from trial-averaged neural data. Delay and response subspaces are determined as the orthogonal subspaces that maximally contain the variance of neural activity in the delay and response epochs, respectively. Correct trials from the DR context were used to identify subspaces. **b.** Activity during DR trials within each subspace for an example session. *Left*, motion energy across trials. *Middle*, sum-squared magnitude of activity in the response subspace. *Right*, sum-squared magnitude of activity in the delay subspace. Trials sorted by average delay epoch motion energy. **c.** Sum-squared magnitude of activity in delay and response subspaces during DR lick-left and lick-right trials for an example session. Mean and standard error across trials shown. **d.** Normalized variance explained of the neural data during DR trials by the activity in the delay and response subspaces during the delay and response epochs. Delay and response subspaces selectively capture activity in the delay and response epochs, as expected. Points indicate sessions, bar height indicates the mean across sessions, and error bars indicate standard deviation across session (*n* = 25 sessions). **e.** Variance explained (R^2^) of motion energy by the sum-squared magnitude of activity in the delay and response subspaces using single-trial DR and WC data. Each point represents the mean across trials for a session (*n* = 25 sessions). **f.** Cross-correlation between motion energy and activity in the delay and response subspaces using single-trial DR and WC data. Lines indicate mean across sessions and shaded region represents standard error of the mean across sessions (*n* = 25 sessions). **g.** Subspace alignment for single units across all sessions (*n* = 483 single units, see **Methods**). Values closer to 1 indicate that more variance of a unit’s activity is contained within the preparatory subspace. **h.** Schematic of the approach for estimating movement-null and movement-potent subspaces from single-trial data. Motion energy is used to annotate when an animal is moving (*pink*) or stationary (*green*). These labels are then applied to single-trial neural data. Movement-null and movement-potent subspaces are determined as the orthogonal subspaces that maximally capture the variance of neural activity during periods of quiescence and movement, respectively. Trials from both DR and WC contexts were used to identify subspaces. **i-n.** Same as (b-f) but using the single-trial approach for estimating movement-null and movement-potent subspaces.

### Subspace decomposition of single-trial neural activity

To avoid these limitations, we modified this analytical approach in two ways. First, because uninstructed movements vary considerably from trial-to-trial, we avoided analyses of trial-averaged data and focused on single-trial data. Second, rather than identifying subspaces capturing variance during the delay (when no movement is assumed) and response epochs, we instead annotated all time points across a session in which animals were moving and all time points at which animals were stationary. This could be straightforwardly achieved using a threshold on motion energy (calculated as the magnitude of frame-by-frame changes in images captured by video). We then found the orthogonal subspaces that best explained variance during all time points across a session at which animals were stationary and moving (**Fig. 5h**). We termed the resulting subspaces ‘movement-null’ and ‘movement-potent’ subspaces. The movement-potent subspace contains dynamical patterns typically associated with movement. Patterns that are observed in the absence of movements – those likely related to cognitive and other ‘internal’ processes – are contained within the movement-null subspace. Following this approach, we found that activity within the two subspaces together explained 72% of the variance in single-trial neural activity (movement-null: 30 ± 9.5%, mean ± s.d.; movement-potent: 42 ± 8.7%; *n* = 25 sessions). Neural dynamics in the movement-potent subspace were correlated with motion energy on single trials, as might be expected (**Fig. 5i,l,m**). Naively, movement-null subspace activity might be expected to be similarly anticorrelated with motion energy because that subspace is constructed to specifically capture variance during periods of animal stationarity. This outcome was largely not observed. Rather, movement-null subspace activity did not display a consistent relationship with movement (**Fig. 5i,l,m**) suggesting that these ‘internal’ dynamics were not only prominent in the absence of movement, but persisted during movement as well, in line with the supposition that cognitive processes can persist during movement.

Uninstructed movements were commonly observed during the delay epoch of DR trials, during motor planning, and were highly correlated with putative choice and urgency signals (**Fig. 3c-h**). We examined whether the movement-potent subspace might then inadvertently capture choice or urgency dynamics that are correlated in time with movements. To examine this possibility, we examined movement-null and movement-potent subspaces estimated using trials from the WC context only. Here, the movement-potent subspace is determined in a context in which choice and urgency signals are absent^23^ (**Fig. 2d** and **EDFig. 4c**), precluding the possibility that dimensions of neural activity related to choice and urgency are inadvertently assigned to the movement-potent subspace due to their correlation with movement. This approach yielded subspaces that were highly similar (**EDFig. 4d-f**). In another control, examining movement-null and movement-potent subspaces estimated only using data recorded during the response epoch of both tasks, when planning and urgency dynamics should be minimal, again yielded similar results (**EDFig. 4g-i**). Finally, we estimated the movement-null subspace as the subspace defined by the top principal components of activity recorded during periods of stationarity and then determined the movement-potent subspace as the top principal components of dynamics not already captured in the movement-null subspace. This latter procedure, which is highly conservative in avoiding the spurious assignment of internal dynamics to the movement-potent subspace, again yielded similar results (**EDFig4. j-l** and **Methods**). Together, these observations suggest that the movement-potent subspace indeed captures movement-related signals, and not dynamics related to internal processes that are somewhat correlated in time with movements.

We hypothesized that activity in the movement-null and movement-potent subspaces should capture variance in neural activity across all task epochs. Animals may think and move simultaneously; thus, it follows that internal dynamics contained in the movement-null space may be present during the response epoch as well. Additionally, uninstructed movements occur during both the sample and delay epochs, concurrent with stimulus and choice encoding. Thus, we should also observe dynamics within the movement-potent subspace during all task epochs. As expected, activity within movement-null and movement-potent subspaces captured variance across epochs (**Fig. 5k**). Interestingly, the proportion of units with activity largely confined to the movement-potent subspace was larger than expected by chance (**Fig. 5n** and **EDFig. 5b**; *p* = 6x10^-19^, *n* = 483 single units, see **Methods**), and larger than when subspaces were identified from trial-averaged data here (**Fig. 5g**) and in non-human primates^39^. This observation suggests the identification of neuronal populations engaged solely in motor processes that could not be identified without properly accounting for uninstructed movements.

One of the few parameters associated with this procedure is the choice of the dimensionality of the movement-null and movement-potent subspaces. We examined whether varying the dimensionality of each subspaces (from four to 13) influenced these results. We found our results to be largely insensitive to subspace dimensionality (**EDFig. 6a-c**), again suggesting minimal sensitivity to choice of parameters.

**Figure 6.**
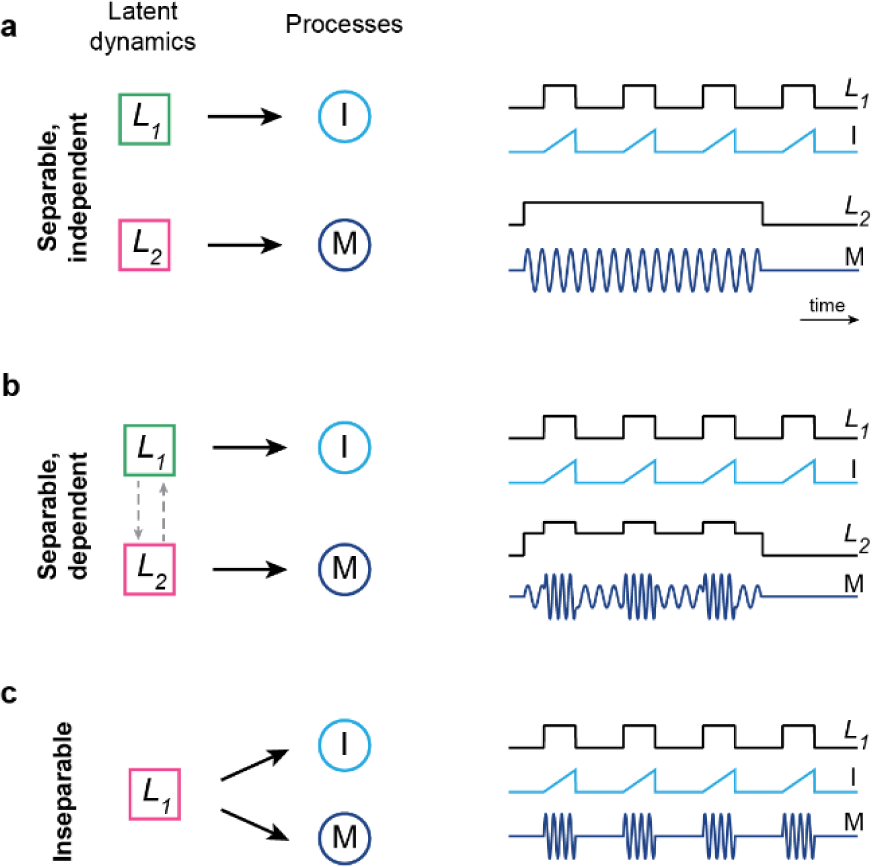
Schematic of potential relationships between internal and movement-related dynamics. **a.** Internal and movement-related dynamics that are independent and separable. *Left,* schematic depicting an internal process (I) and a motor process (M) that are each governed by separate latent dynamics, *L_1_* and *L_2_*, respectively. *L_1_* and *L_2_* evolve within the movement-null (green) and movement-potent (pink) subspaces, respectively. *Right,* cartoon time series of separable, independent latent dynamics (*L_1_* and *L_2_*) and related processes (I and M). **b.** Same as (a) in the case of latent dynamics which are separable but dependent. I and M may be loosely correlated in time. **c.** Same as (a) and (b) in the case of inseparable processes governed by a single set of latent dynamics (*L_1_*).

### Internal and movement-related dynamics during motor planning

The movement-potent subspace may contain at least three classes of movement-related neural dynamics: (1) motor commands (and efference copies), (2) sensory feedback related to movements, and (3) internal dynamics that are present *exclusively during periods of* movement. That is, if a particular internal process is “embodied,” in the sense that it is mediated by the same latent dynamics as responsible for associated movements – and thus always observed in the presence of those movements - then we would expect to find those latent dynamics wholly contained within the movement-potent subspace (**Fig. 6c**). Dynamics supporting internal processes which are independent of movements should be contained within the movement-null subspace (**Fig. 6a**). Between these extremes, we expect to observe dynamics within both subspaces when a particular internal process (with associated latent dynamics in the movement-null subspace) biases the expression of movements (with associated latent dynamics in the movement-potent subspace) (**Fig. 6b**). In that latter case, internal processes may appear correlated with movements despite being largely mediated by distinct latent dynamics.

To evaluate these possibilities, we examined whether putative cognitive signals (illustrated in **Fig. 2**) evolved within the movement-potent and/or movement-null subspaces. We found choice selectivity – dynamics that differentiate left and right DR trials – in both subspaces. The existence of choice dynamics in the movement-null subspace indicates an internal representation of choice that is separable from dynamics related to the execution of choice-related uninstructed movements (see **Fig. 6b**). The average time courses of choice dynamics in these subspaces displayed subtle, yet notable, differences, as did their expression on error trials. Selectivity emerged quickly following stimulus presentation in the movement-null subspace and did not change significantly through the delay epoch (**Fig. 7a**; *p* > 0.05, paired *t*-test comparing selectivity during last 100ms of sample and delay epochs, *n* = 25 sessions **EDFig. 7c**) consistent with an internal representation driven by sensory input. On error trials, selectivity in the movement-null subspace initially followed the same trajectory as correct trials but decayed following stimulus offset. In the movement-potent subspace, in contrast, selectivity increased slowly during the sample epoch and continued to increase monotonically during the delay epoch (*p* = 3x10^-13^, paired *t*-test comparing selectivity during last 100ms of sample and delay epochs, *n* = 25 sessions; *p* = 1x10^-5^ comparing change in selectivity during the delay epoch in movement-potent and movement-null subspaces; **EDFig. 7b**), and no significant movement-potent subspace selectivity emerged on error trials. These observations suggest that sensory stimuli initially drive appropriate dynamics within the movement-null subspace on error trials, but that choice is not encoded stably and does not engage movement-potent dynamics related to uninstructed movements.

**Figure 7.**
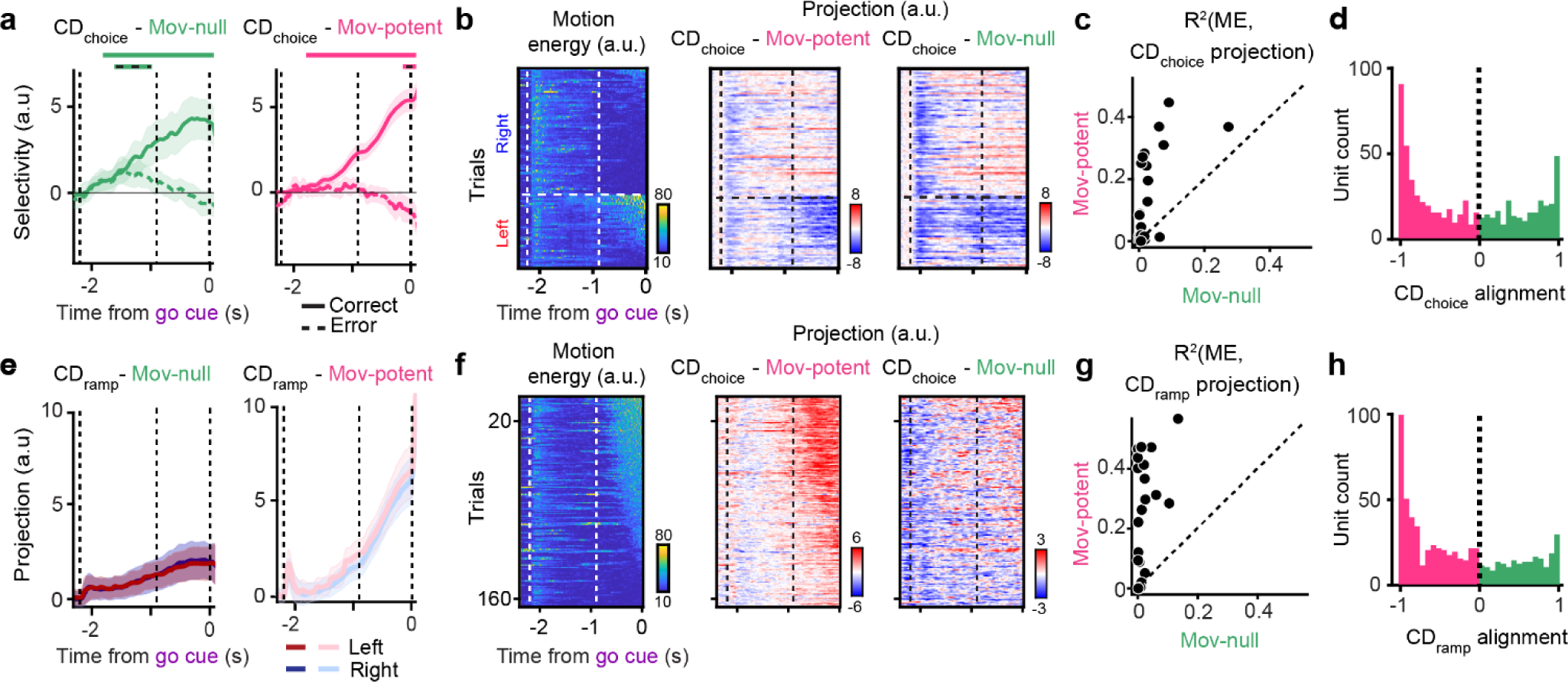
Subspace decomposition allows for the re-examination of population measures of motor planning. **a.** Selectivity (neural projections on lick-right trials minus lick-left trials) of movement-null (*left*) and movement-potent (*right*) subspace activity when projected along CD_choice_. Mean and 5-95% CI of the bootstrap distribution for correct (solid) and error (dashed) trials shown. Horizontal bars (solid lines - correct trials; solid lines with black dashes - incorrect trials) above plot indicate when the selectivity trace is significantly different from zero (*p* < 0.05, one-sided test, bootstrap). **b.** Projections along CD_choice_ in the movement-potent subspace (*middle*), but not the movement-null subspace (*right*), follow the time-course and magnitude of motion energy (*left*) on single lick-left and lick-right trials (example session). **c.** Variance explained (R^2^) of motion energy by projections along CD_choice_ in the movement-null or movement-potent subspace on single trials. Each point is the average across all trials in a session. **d.** Distribution of single unit alignment with CD_choice_ (*n* = 483 single units, see **Methods**). Distribution of movement-potent tuned units (alignment ≤ -0.8) was significantly different than expected by chance (*p* = 1x10^-13^, see **Methods**). Similarly, distribution of movement-null tuned units (alignment ≥ 0.8) was significantly different than expected by chance (*p* = 7x10^-12^). **e**. Projection of movement-null (*left*) and movement-potent (*right*) subspace activity along CD_ramp_ on lick-left and lick-right trials. Mean and 95% CI of the bootstrap distribution for correct trials shown. **f-h**. Same as (b-d) for CD_ramp_. **h.** Distribution of single unit alignment with CD_ramp_ (*n* = 483 single units). Distribution of movement-potent tuned units (alignment ≤ -0.8) was significantly different than expected by chance (*p* = 6x10^-14^). Similarly, distribution of movement-null tuned units (alignment ≥ 0.8) was significantly different than expected by chance (*p* = 1x10^-12^).

Although the trajectories of choice dynamics in each subspace shared some similarities in trial-averaged data (**Fig. 7a**), the existence of choice dynamics in both subspaces implies that they must differ markedly on single trials. The onset and magnitude of activity along the component of CD_choice_ within the movement-potent subspace tracked trial-type selective motion energy on a moment-by-moment basis (**Fig. 7b, c**), as expected, while there was no consistent relationship between motion energy and activity along the component of CD_choice_ in the movement-null subspace. Following the go cue, transient responses accompanying movement initiation were absent from the movement-null subspace (**EDFig. 7a**), further suggesting the existence of choice dynamics which are separable from ongoing movement dynamics. Importantly, we found that many single units contribute to either the movement-null or movement-potent subspace representations of CD_choice_, but not both (**Fig. 7d** and **EDFig. 6d**), providing additional evidence of dynamics that are not only separable, but encoded by distinct populations of neurons (**Fig. 7d**) within the ALM microcircuit.

In contrast, we found that ramping dynamics were mostly confined to the movement-potent subspace (**Fig. 7e**) – perhaps surprising given that ramping activity has been interpreted as an internal urgency or timing signal^69–72^. This observation remained consistent when estimating movement-null and movement-potent subspaces using only WC trials or using only the response epoch of WC and DR trials (**EDFig. 8a**), when ramping dynamics were absent, confirming that ramping dynamics were not observed in the movement-potent space because of the mis-assignment of internal ramping dimensions to the movement-potent subspace because of their correlation in time with movement. We further searched for the existence of movement-null subspace ramping dynamics by explicitly identifying the dimension within the movement-null and movement-potent subspaces that maximized ramping dynamics – which could be different than the dimension that maximized ramping within the full activity space – and again failed to identify a prominent ramping signal in the movement-null subspace (**EDFig. 8b**). The relative paucity of ramping dynamics in the movement-null subspace suggests that they are not readily separable from associated movements in the ALM in our behavioral paradigm (see **Discussion**).

**Figure 8.**
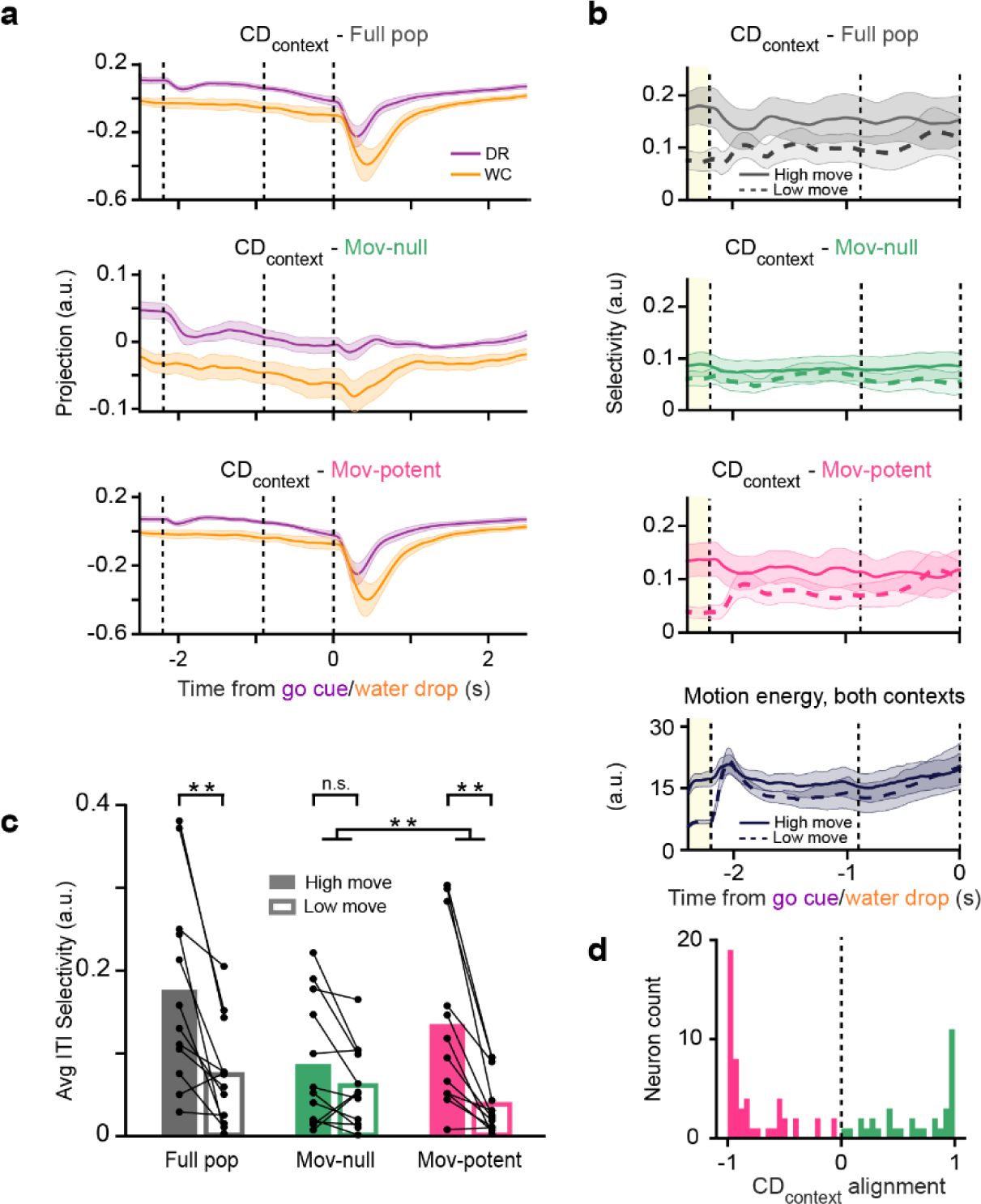
A persistent, cognitive representation of context in the movement-null subspace. **a.** Projections along CD_context_ on DR hit trials (*purple*) and WC hit trials (*orange*) within the full population (*top*), movement-null subspace (*middle*), or the movement-potent subspace (*bottom*). Solid lines denote the mean projection across sessions (*n* = 12 sessions). Note that transient response epoch dynamics are absent in the movement-null subspace. **b.** *Top three panels*, selectivity in CD_context_ projections (DR – WC trials) on ‘high move’ (*solid lines*) or ‘low move’ (*dashed*) trials. Lines denote the mean selectivity across sessions (n = 12) in the full space of neural activity (*top*), activity within the movement-null subspace (*second*), and movement-potent subspace (*third*); *Bottom panel*, average motion energy across sessions on ‘High move’ or ‘Low move’ trials. DR and WC trials are grouped together for each session. Shaded area, 95% CI across sessions. Yellow region denotes ITI used to define ‘High move’ and ‘Low move’ trials. **c.** Average CD_context_ selectivity during the ITI (yellow shaded region in (b)) for ‘High move’ (*filled bars*) vs. ‘Low move’ (*open bars*) trials. Dots denote single sessions; bars, mean across sessions (*n* = 12 sessions). Asterisks denote significant differences (**, *p* < 0.01) in CD_context_ selectivity (*p*-values from left to right comparing ‘High move’ vs. ‘Low move’ trials, Full population: *p* = 0.003, movement-null: *p* = 0.095, movement-potent: *p* = 1.6x10^-3^; Δselectivity between ‘High move’ and ‘Low move’ trials in movement-null vs. movement-potent: *p* = 9.5x10^-3^, paired *t*-test). **d.** Distribution of single unit alignment with CD_context_ (see **Methods**). Distribution of movement-potent tuned units (alignment ≤ -0.8) was significantly different than expected by chance (*p* = 2x10^-8^, see **Methods**). Similarly, distribution of movement-null tuned units (alignment ≥ 0.8) was significantly different than expected by chance (*p* = 9x10^-6^, *n* = 84 single units). Lines with shaded regions depict mean and 95% confidence intervals across sessions throughout.

### A persistent movement-null subspace representation of context

Next, we examined projections of activity along CD_context_ in each subspace (**Fig. 8a**). We found robust contextual selectivity in both subspaces, consistent with the interpretation that the ALM contains both an internal representation of context and activity related to context-dependent movements. The transient, response-epoch dynamics observed along CD_context_ (**Fig 2d**, *far right*) were entirely contained within the movement-potent subspace (**Fig. 8a**) likely indicative of subtle context-specific differences in instructed movements. We then compared context selectivity (the difference in projections onto CD_context_ on DR vs. WC trials) on trials with high motion energy during the ITI (‘High move’ trials) and trials with little or no ITI motion energy (‘Low move’ trials). Context selectivity within the movement-null subspace was indistinguishable in ‘High move’ and ‘Low move’ trials. In contrast, context selectivity within the movement-potent subspace was reduced by 72% on ‘Low move’ trials (**Figs. 8b,c**; Full population: selectivity reduced by 0.101 ± 0.092, mean ± s.d., *p* = 0.003; movement*-*null: 0.026 ± 0.050, *p* = 0.095; movement*-*potent: 0.098 ± 0.082, *p* = 1.6x10^-3^, paired *t*-test; Δ selectivity between ‘High move’ and ‘Low move’ trials in movement-null vs. movement-potent: *p* = 9.5x10^-3^, paired *t*-test). The magnitude and timing of the reduction in movement-potent subspace context selectivity mirrored the reduction in motion energy on ‘Low move’ trials (**Fig. 8b***, bottom*). Together, these observations demonstrate that the movement-null – but not movement-potent – subspace contains a stable representation of context that is unchanged in the presence of both instructed and uninstructed movements. Further, we found that largely distinct populations of single neurons contribute preferentially to the movement-null and movement-potent representations of context (**Fig. 8d**). These observations suggest that the analysis of context-dependent dynamics without subspace decomposition indeed spuriously conflated separate latent dynamics, encoded by different populations of neurons, likely responsible for internal representations of context (contained within the movement-null subspace) and related context-dependent movements (contained in the movement-potent subspace).

### Encoding of tongue kinematics in the movement-potent subspace

Finally, we asked whether trial-type dependent activity following the go cue relates to movement, according to previous suppositions^58,6458,64^ (**EDFig. 10**). Left-right selectivity along CD_action_ existed within both the movement-null and movement-potent subspaces (**EDFig. 10a**), although the magnitude of selectivity in the movement-null subspace was substantially smaller in magnitude. Interestingly, on error trials, activity along CD_action_ flipped to resemble that of the other trial type in the movement-null subspace but flipped asymmetrically within the movement-potent subspace. This asymmetry closely corresponded to an asymmetry in tongue angle, with incorrect movements directed to less extreme angles when directed to the left, on average (**EDFig. 10b,c**). The component of CD_action_ within the movement-potent subspace also better tracked moment-to-moment changes in tongue angle during the response epoch (**EDFig. 10d,e**; *p* = 2 x10^-8^, paired *t*-test between movement-null and movement-potent variance explained of tongue angle, *n* = 25 sessions). The similarity between tongue angle, a kinematic feature, and dynamics only within the movement-potent representation of CD_action_ is notable, as our analytical approach to the identification of subspaces does not incorporate any kinematic information. The interpretation of action dynamics within the movement-null subspace remains unclear but could relate to an internal representation of the motor plan, or intention of the animal, to respond to one reward port or the other.

## Discussion

We aimed to understand whether the neural correlates of three cognitive variables commonly examined in sensorimotor decision-making tasks – choice, urgency, and context – were separable from the neural encoding of related uninstructed movements in the mouse ALM. We addressed this question by adapting an analytical formalism in which neural activity is decomposed into orthogonal subspaces – here, a movement-potent subspace containing dynamics related to the execution of movements and a complementary movement-null subspace containing dynamics related to cognitive and other internal processes^39,40,50^. Extending upon this framework allowed us to consider single-trial neural data recorded in the presence of uninstructed movements that can vary dramatically across trials in their timing and identity (**Fig. 5**).

### Separability of choice, urgency, and context encoding from movement-dynamics

Using this approach, we demonstrated that the ALM contains neural dynamics related to encoding of choice and context that could be readily separated from movements, despite both cognitive variables being strongly correlated with movements in time (**Fig. 7** and **Fig. 8**). Choice-selective signals were present in both movement-null and movement-potent subspaces (**Fig. 7a**), consistent with the interpretation that a cognitive representation of choice within the movement-null subspace biases the probability and identity of uninstructed movements in a choice-selective manner. Supporting this interpretation, movement-null and movement-potent subspace dynamics followed similar trajectories, on average, but differed on single trials (**Fig. 7b**). Further, movement-null subspace encoding of choice exhibited features absent from encoding within the movement-potent subspace. Choice encoding increased in the presence of external stimuli, while movement-potent subspace encoding continued to increase monotonically after sensory stimuli were removed, mirroring the temporal evolution of animal’s uninstructed movements. On error trials, stimulus-driven choice encoding initially evolved correctly – but only within the movement-null subspace – before decaying prior to the go cue (**Fig. 7a**). Together, these results suggest choice-related cognitive and motor processes that are governed by separable latent dynamics (**Fig. 6b**).

Context was stably encoded within the movement-null subspace during all trial epochs and the inter-trial interval (**Fig. 8a-c**). Although context was strongly represented in the movement-potent subspace as well, these dynamics were significantly reduced in the absence of movement and were also modulated by animals’ instructed movements. These results suggest a persistent internal representation of context in the movement-null subspace in addition to distinct movement-potent-subspace dynamics related to context-dependent movements, again indicative of related, but separable, latent dynamics (**Fig. 6b**).

Surprisingly, ramping dynamics proposed to underlie urgency (or timing)^69–72^ were principally represented in the movement-potent subspace (**Fig. 7e**) indicating the possible absence of an internal representation of urgency that can be dissociated from movements in the ALM. Although it remains possible that a separable representation of urgency exists elsewhere in the brain, our results could alternatively imply that urgency is typically embodied, or inseparable from movement, in our behavioral paradigm^74,75^.

In contrast to these cognitive variables, encoding of kinematic features of movements, which were not used to determine subspaces, were confined to the movement-potent subspace (**EDFig. 10**), consistent with our interpretation of these subspaces.

### Subspace decomposition

The approach to subspace decomposition presented here represents a means for assessing the issue of separability, and for isolating separable dynamics into movement-related and internal components for further analyses across a wide range of experimental preparations. This method only requires annotation of when animals are moving and utilizes a robust analytical approach that does not require fine-tuning of parameters.

A number of alternative methods have been proposed to evaluate neural dynamics associated with cognitive processes in the presence of movement. The most common approach, which assumes separability of cognitive and motor dynamics, is to parameterize movements as fully as possible and ‘regress them out.’ However, parameterization of movement, particularly the orofacial and postural movements that have been associated with strong, brain-wide neural dynamics^30,31,76^, can be challenging. Further, this approach typically assumes a linear relationship between neural activity and kinematic (or electromyographic) variables – an assumption unlikely to hold for common movements mediated by central pattern generators and other highly nonlinear mechanisms^77–79^. Subspace decomposition is straightforward to implement under a range of experimental conditions and is entirely consistent with complex, nonlinear relationships between neural activity and movement variables, as expected for many orofacial movements^77^ and locomotion. Methods for properly interpreting cognitive signals in the presence of related movements on single trials will be vital for increasingly common experimental paradigms examining freely moving animals performing complex, naturalistic behaviors^80–90^.

Determination of movement-null and movement-potent subspaces as the orthogonal subspaces containing the most variance in neural activity during periods of stationarity and movement is conservative in the assignment of dynamics to the movement-null subspace. It is more likely that dynamics associated with cognitive and other internal processes are misassigned to the movement-potent subspace than vice-versa. For example, dynamics associated with “embodied” cognitive processes that always occur during periods of movement will only be represented in the movement-potent subspace (**Fig. 6c**). Thus, the dynamics within the movement-null subspace are highly likely to indicate signals related to internal processes. Straightforward variations of this approach can be used to determine subspaces in a manner that is more conservative in assigning dynamics to the movement-potent subspace – for example, estimating subspaces from response epoch data (‘instructed’ movements), from WC trials (where uninstructed movements occur in the absence of choice and urgency signals) or simply through principal component decomposition of activity recorded during periods of stationarity (**EDFig. 4j-l**). Nevertheless, the similarity of results obtained using these analytical variations argue against the possibility that cognitive dynamics associated with choice and urgency were inadvertently assigned to the movement-potent subspace, due to their correlation in time with uninstructed movements, in this study.

That many individual units contribute solely to one subspace (**Figs. 5n, 7d,h, 8d**) suggests that the complementary subspaces we identify could map to distinct cell types within the underlying cortical circuit^63,91^. The suggestion that individual neurons only appear to code for either internal or movement-related variables when uninstructed movements are accounted for (**Figs. 5n, 7d,h, 8d**) underscores the importance of properly considering movements in future work focused on the roles of functionally, anatomically, and/or transcriptomically-defined cell-types in neural computation.

Our approach to subspace decomposition makes several simplifying assumptions. First, movements of the posterior torso, hindlimbs, or tail of the animal were not routinely captured via videography. It remains possible, albeit unlikely, that animals routinely perform movements confined to these portions of the body without concomitant movement of the forepaws, head, neck, face, or whiskers^26^. Periods of stationarity were no doubt algorithmically classified imperfectly and could have contained subtle movements below our threshold for detection. Second, we did not consider potential time lags between motion energy captured via videography and the dynamics associated with motor and/or sensory signals^73^. However, these time lags, on the order of a few tens of milliseconds, should be much shorter than the timescale of dynamics explored in this study. Third, the set of signals related to movement and/or cognitive processes may also be better described as occupying nonlinear manifolds rather than linear subspaces^92,93^. Subspaces (or manifolds) could also shift dynamically during behavior following changes in the activation of upstream or downstream brain regions^94,95^ as a result of changes in context^16,48,96^. Considering this additional complexity may improve estimates of the dynamics associated with specific neural processes.

### Variable relationships between cognition and movement

The dynamics associated with some cognitive processes and movements may be largely independent (**Fig. 6a**) – perhaps in the case of the locomotor patterns of an individual walking down the street while thinking about what they will cook for dinner. On the other hand, some cognitive processes may be embodied, in the sense that they are tightly linked to externally observable changes in the body (**Fig. 6c**), such as the relationship between arousal and changes in pupil diameter^97–99^. Between these extremes may be movements which are correlated with - but mediated by neural dynamics that are separable from - those supporting cognitive and other internal processes. The probability and identity of uninstructed movements in this study appeared related to choice and context (**Fig. 6b**), but with a high degree of trial-to-trial and moment-to-moment variability unlikely to reflect commensurate variability in the cognitive processes to which they relate. Correlations between cognitive processes and movements may also differ considerably in trained, head-fixed animals compared to naturalistic settings^83,88^. This high degree of variability in these relationships underscores the importance of developing and utilizing tools for assessing whether the neural activity supporting cognitive processes can be dissociated from those related to movements in particular experimental paradigms of interest.

The tight link between cognition and movement might suggest that some cognition-associated movements may be beneficial for behavior. For example, postural adjustments may be highly related to decisions or motor plans, enabling faster reaction times and/or more accurate movements^76,100^. Movements are also essential for some internal processes – those supporting active sensation of the environment surely facilitate the construction and continual updating of internal models of the environment^30,46^. However, just as separability may not be consistent across movements and cognitive processes, not all cognition-associated movements may have a functional role^30,101^. A poker player’s ‘tell’ may be highly related to their cognitive state, yet may not directly aid the player in the game. Regardless of their functional relationship, understanding the separability of cognitive processes and associated movements in a wide variety of experimental settings is essential for the interpretation of neural dynamics observed during behavior.

## Methods

### Animals

This study used data collected from 17 mice; both male and female animals older than 8 weeks were used. Six animals were used for the two-context task and were either C57BL/6J (JAX 000664) or VGAT-ChR2-EYFP (-/-). An additional three animals, either C57BL/6J or VGAT-ChR2-EYFP (-/-) were trained only on the delayed response task. A separate cohort of four animals were used in the randomized delay task (VGAT-ChR2-EYFP (+/-) or VGAT-ChR2-EYFP (-/-)). Finally, four VGAT-ChR2-EYFP (+/-) animals were used in optogenetics experiments. Mice were housed in a 12-hour reverse dark/light cycle room with ad libitum access to food. Access to water was restricted during behavioral and electrophysiology experiments (see ***Mouse behavior***). Sample sizes were not determined using any statistical tests.

### Surgical procedures

All surgical procedures were performed in accordance with protocols approved by the Boston University Institutional Animal Care and Use Committee. For post-operative analgesia, mice were given ketoprofen (0.1 mL of 1 mg/mL solution) and buprenorphine (0.06 mL of 0.03 mg/mL solution) prior to the start of all surgical procedures. Mice were anesthetized with 1-2% isoflurane and placed on a heating pad in a stereotaxic apparatus. Artificial tears (Akorn Sodium Chloride 5% Opthalmic Ointment) were applied to their eyes and a local anesthetic was injected under the skin (Bupivacaine; 0.1 mL of 5 mg/mL solution) above the skull. The skin overlying the skull was removed to expose the ALM (AP: +2.5 mm, ML: +1.5 mm), bregma, and lambda. The periosteum was removed and the skin was secured to the skull with cyanoacrylate (Krazy Glue) around the margins. For electrophysiology experiments, a headbar was implanted just posterior to bregma and secured with superglue and dental cement (Jet™). Wells to hold cortex buffer (NaCl 125mM, KCl 5mM, Glucose 10mM, HEPES 10mM, CaCl2 2mM, MgSO4 2mM, pH 7.4) during electrophysiology recordings were sculpted using dental cement and a thin layer of superglue was applied over any exposed bone.

For optogenetics experiments, after headbar implantation, bone overlying frontal cortex was thinned bilaterally with a dental drill and removed, exposing the frontal cortex. A glass window was then implanted over each hemisphere and secured to the skull with cyanoacrylate^102^. Before performing photoinactivation experiments, any bone regrowth was removed.

### Mouse behavior

Following surgery, mice were allowed to recover for ∼1 week and then water restricted, receiving 1.0 mL of water per day. Behavioral training commenced after animals had been water restricted for 3-4 days. If animals received less than 1.0 mL during training, they were supplemented with additional water.

All mice used in experiments were first trained on the delayed-response task with a fixed delay epoch until they reached at least 70% accuracy. At the beginning of each trial, one of two auditory tones lasting 1.3 s were played; the tone indicating a ‘right’ trial was white noise while the tone indicating a ‘left’ trial was a pure tone (8 kHz pulses). The delay epoch (0.9 s for 12 mice, 0.7 s for 1 mouse and linearly time warped to 0.9 s) started after the completion of the sample tone. Following the delay epoch, an auditory go cue was played (swept-frequency cosine [chirp], 10 ms) after which animals were rewarded with ∼3 µL of water for contacting the correct lickport. Lickport contacts before the response epoch resulted in the current task epoch restarting, ensuring that they could not progress to the next task epoch until they had stopped licking for the specified length of the current epoch. If the animal did not respond within 3 s of the go cue, this was considered an ‘ignore’ trial, although responses were typically registered within 300 ms of the go cue.

Mice used for the two-context task (**Figs. 1,2,4,8** and **EDFig. 1-3**) were introduced to the water-cued context after they were fully trained on the delayed-response context and at least two days before the first electrophysiology recording session. A behavioral session began with ∼100 DR trials and was then followed with alternating blocks of WC and DR trials. Each interleaved block was 10-25 trials, with no fixed block duration. All sessions started with DR trials. Pilot sessions beginning with WC trials had to be terminated early due to a high ‘no response’ rate, likely due to the mouse becoming sated early in the session. In a WC trial, all auditory cues were omitted and a ∼3 µL water reward was presented randomly from either lickport. Trials in which the animal contacted the lickport prior to the reward (‘early lick’) were omitted from analyses. Inter-trial intervals were randomly drawn from an exponential distribution with mean 1.5 s. There were no explicit cues during the ITI to indicate to the animal which context block it was in.

For the randomized delay task (**Fig. 3g-h**), mice were first fully trained on the fixed delay DR task (delay duration of 0.9 s). The delay epoch length was then randomized – the duration was randomly selected from six possible values (0.3, 0.6, 1.2, 1.8, 2.4, and 3.6 s). The probability of each delay duration being selected was assigned such that it resembled a probability density function of an exponential distribution with τ = 0.9 s, as in ref ^65^. Animals were trained on this version of the task until they became experts (> 70% accuracy and < 20% early lick rate).

### Videography analysis

High-speed video was captured (400 Hz frame rate) from two cameras (FLIR, Blackfly® monochrome camera, Model number: BFS-U3-16S2M-CS). One provided a side view of the mouse and the other provided a bottom view. We tracked the movements of specific body features using DeepLabCut^52^ (**Figs. 1,3,4**). The tongue, jaw, and nose were tracked using both cameras while the paws were only tracked using the bottom view. Position and velocity of each tracked feature was calculated from each camera. The x and y position of each kinematic feature was extracted from the output of DeepLabCut. Missing values were filled in with the nearest available value for all features except for the tongue. The velocity of each feature was then calculated as the first-order derivative of the position vector. Tongue angle and length were found using the bottom camera. Tongue angle was defined as the angle between the vector pointing from the jaw to the tip of the tongue and the vector defining the direction the mouse was facing. Tongue length was calculated as the Euclidean distance from the jaw to the tip of the tongue.

Plots of kinematic feature overlays (**Figs. 1e, 3a, 4a** and **EDFig. 3**) were generated by plotting an [r, g, b] value for each timepoint, *t*, where the values were specified by [KinFeature1_t_, KinFeature2_t_, KinFeature3_t_]. All kinematic features (speed or motion energy) were first standardized by taking the 99^th^ percentile across time and trials and normalizing to this value.

### Motion energy

The motion energy for a given frame and pixel was defined as the absolute value of the difference between the median value of the pixel across the next 5 frames (12.5 ms) and the median value of the pixel across the previous 5 frames (12.5 ms). Motion energy for each frame was then converted to a single value by taking the 99^th^ percentile (∼700 pixels) of motion energy values across the frame. Motion energy calculated in this manner was high during overt movements over small regions of pixel space, such as during whisking, while remaining relatively low during passive respiration that corresponded to subtle motion across the animal’s body. A threshold above which an animal was classed as moving, was defined on a per session basis manually. Motion energy distributions, per session, were bimodal, showing a large peak with little variance at low motion energy values and a second, smaller peak, with high variance at large values. The threshold was set as the motion energy value separating these two modes. We found that this method of setting the threshold captured both short and long duration movements. Alternative methods, such as defining a baseline period of no movement against which to compare, were less successful due to the variability in timing of the movements across trials.

### Photoinactivation experiments

Optogenetic photoinactivation was deployed on ∼30% of trials selected at random. A ‘masking flash’ (470 mm LED’s LUXEON REBEL LED) controlled by an Arduino Teensy microcontroller (100 ms pulses, at 10 Hz) was delivered constantly for the duration of the session to prevent mice from differentiating control and photoinactivation trials. A 488-nm laser (Coherent OBIS 488 nm LX 150 mW Laser System) was used for all photoinactivation experiments. ChR2-assisted photoinactivation (**Fig. 1b,c,g-** and **EDFig. 1b-d**) was performed through a cranial window^102^ (see ***Surgical Procedures***), which replaced bone over the frontal cortex. Light was delivered either at the start of the delay epoch, or at the start of the response epoch (only one epoch tested per session). A power density of 1.5 mW/mm^2^ was used for all photoinactivation experiments.

For delay epoch photoinactivation (**Fig. 1g-i** and **EDFig. 1b**), light was delivered at the onset of the delay epoch and lasted for 0.6 s followed by a 0.2 s linear ramp down. We targeted one of three regions per session: bilateral motor cortex (ALM and tjM1), bilateral ALM, or bilateral tjM1. A scanning galvo system (THORLABS GPS011) was used to simultaneously target both hemispheres by scanning at 40 Hz. The beam (2 mm diameter) was centered around the following locations: ALM (AP 2.5 mm, ML 1.5 mm), tjM1 (AP 1.75 mm, ML 2.25 mm), motor cortex (i.e. ALM and tjM1; AP 2.25 mm, ML 1.85 mm). Due to their proximity, when specifically targeting ALM or tjM1, Kwik-Cast™ (World Precision Instruments) was applied to the surrounding regions to prevent photoinactivation of other regions. For photoinactivation at the go cue (**Fig. 1b,c** and **EDFig. 1c,d**), light was delivered to the motor cortex for 0.8 s followed by a 0.2 s ramp down beginning at the go cue presentation.

### Electrophysiology recordings

Extracellular recordings were performed in the ALM using one of two types of silicon probes: H2 probes, which have two shanks, each with 32 channels with 25-µm spacing (Cambridge Neurotech) or Neuropixels 1.0^103^ which have one shank and allow recording from 384 channels arranged in a checkerboard pattern (IMEC). For recordings with H2 probes, the 64-channel voltage signals were multiplexed using a Whisper recording system (Neural Circuits, LLC), recorded on a PXI-8133 board (National Instruments) and digitized at 14 bits. The signals were demultiplexed into 64 voltage traces sampled at 25 kHz and stored for offline analysis.

At least 6 hours before recording, a small craniotomy (1-1.5 mm diameter) was made over ALM (AP 2.5 mm, ML 1.5 mm). 2-5 recordings were performed on a single craniotomy on consecutive days. After inserting the probes between 900 and 1100 µm (MPM System, New Scale Technologies), brain tissue was allowed to settle for at least 5 minutes before starting recordings. All recordings were made using SpikeGLX (https://github.com/billkarsh/SpikeGLX).

### Behavioral analysis

All sessions used for behavioral analysis had at least 40 correct DR trials for each direction (left or right) and 20 correct WC trials for each direction, excluding early lick and ignore trials, which were omitted from all analyses. To assess the impact of ALM photoinactivation on movement initiation, we first calculated the percent of trials with a correct lick within 600 ms of the go cue/water drop (**Fig. 1c**). To find the fraction of time with the tongue visible during the photoinactivation period (**EDFig. 1d**), for each trial we found the number of time-points within the 1 s after the go cue/water drop where the tongue was labelled as visible by DeepLabCut and divided that by the total number of time-points within the photoinactivation period. This was then averaged for all control or photoinactivation trials for a given session.

To assess the impact of bilateral MC/ALM/tjM1 photoinactivation during the delay epoch, the average motion energy during the delay epoch was calculated separately for right and left control vs. photoinactivation trials (**Fig. 1h** and **EDFig. 1b**).

### Electrophysiology recording analysis

JRCLUST^104^ (https://github.com/JaneliaSciComp/JRCLUST) and/or Kilosort3^105^ (https://github.com/MouseLand/Kilosort) with manual curation using Phy (https://github.com/cortex-lab/phy) were used for spike sorting. A unit was considered a well-isolated single unit based on manual inspection of its ISI histogram, separation from other units, and its stationarity across the session^106^. Units that passed manual curation but had a higher ISI violation rate were called multiunits. Recording sessions were only included for analysis if they had at least 10 units (see **EDFig. 2** for a distribution of neuron counts across sessions). For the DR task, we recorded 1651 units (483 single units) in ALM from 25 sessions using 9 mice. In 12 sessions from 6 mice, animals performed the two-context task. 522 units (214 well-isolated single units) were recorded in these sessions. Finally, for the randomized delay task, we recorded 845 units (288 well-isolated single units) in ALM from 19 sessions using 4 mice. For subspace alignment (**Figs. 5g,n**, **7d,h**, and **Fig. 8d**) and single-unit selectivity analyses (**Fig. 3g** and **EDFig. 4c**) only well-isolated single units with firing rates exceeding 1 Hz were included. All units with firing rates exceeding 1 Hz were included in all other analyses.

To find choice-selective neurons, forty trials were subsampled for both right and left correct trials and a *ranksum* test was used to compare spike counts for each unit during the sample, delay, or response epochs. Selective units were those with significantly different spike counts (*p* < 0.05) during a given epoch. Context-selective units were defined in a two-step process to ensure that we were not conflating context selectivity with slow changes in animal engagement/motivation over each session. First, forty trials each were subsampled for DR and WC trials and spike counts were compared (*p* < 0.05, *ranksum* test) during the ITI (the 300 ms preceding the sample tone onset) when the animal received no external cues to indicate which context it was in. Because sessions always began with a DR block and often ended with a WC block, it is possible that differences in firing rates across contexts defined in this way could be representing time within the session. To account for this, for each unit identified in the first step, we calculated the difference in spike rate for each pair of DR and WC blocks (e.g. a session with five blocks, DR-WC-DR-WC-DR, would have nPairs=4 adjacent, overlapping block pairs) and included units as context-selective only if their *preferred* context (context with a higher spike rate) was the same for at least nPairs - 1 of the block pairs.

Selectivity in the neural population (**EDFig. 4c**) was calculated as the difference in spike rate on *preferred – non-preferred* trials in choice-selective units, with *preferred* trials referring to the trial type with a higher spike rate for each unit.

### Coding direction analysis

Coding directions (CD) were defined as directions in neural activity state space, defined by firing rates, that maximally separated trajectories of two conditions.

CD_choice_ and CD_action_ were calculated as:

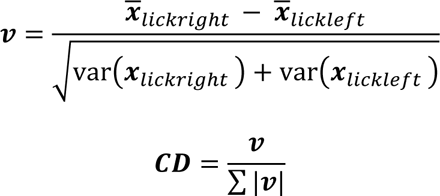

For each unit, the mean spike rate difference between right lick trials, ***x̄***_*lickright*_, and left lick trials, ***x̄***_*lickleft*_, was calculated in a 400 ms time interval. ***x̄***_*lickright*_ and ***x̄***_*lickleft*_ were then individually baseline subtracted and scaled by baseline standard deviation before calculating CDs (baseline: -2.4 to -2.2 seconds relative to the go cue, during the ITI). CD_choice_ was calculated in the last 400 ms of the delay epoch and CD_action_ was calculated in the first 400 ms of the response epoch. The vector ***ν*** was then obtained by normalizing by the square root of the sum of the variances across conditions. Finally, ***ν*** was normalized by its L_1_ norm to ensure projections do not scale with the length of ***ν***, the number of units simultaneously recorded.

CD_ramp_ was calculated as:

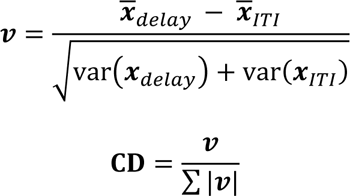

This calculation was similar to the calculations of CD_choice_ and CD_action_but incorporated data from all correct DR trials during the last 400 ms of the delay epoch, ***x̄***_*delay*_, and during the ITI (300 ms preceding the sample tone onset), ***x̄***_***ITI***_. ***x̄***_*delay*_ and ***x̄***_***ITI***_ were both standardized using the baseline statistics from all correct DR trials.

To find CD_context_, we first calculated a CD_context_^*p*^ for each pair, *p*, of DR and WC blocks in a session. If the session ended with a WC block, that block was excluded. CD_context_^*p*^ was calculated as:

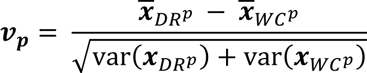

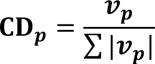

CD_context_ was then defined as the average over all CD_context_^*p*^ in a session. The calculation of CD_context_^*p*^ was similar to the calculation of other CDs but incorporated data from the ITI of correct DR, ***x̄***_*DR^p^*_, and correct WC, ***x̄***_*WC^p^*_, trials in a given pair of blocks. ***x̄***_*DR^p^*_ and ***x̄***_*WC^p^*_ were standardized using the baseline statistics from all correct DR and WC trials.

CD_action_ was orthogonalized to CD_choice_ to exclusively capture response epoch selectivity that emerges after the go cue. CD_ramp_ was orthogonalized to CD_action_ and CD_choice_ to remove selectivity that may emerge during the response epoch that is also captured in CD_action_. Orthogonalization was performed using the Gram-Schmidt process.

Projections of the population activity, ***x*** ∈ ℝ^(B*K)^ ^×^ ^N^ along the CD were calculated as:

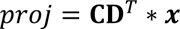

where B is the number of time bins, K the number of trials, N the number of neurons, and *T* is the transpose operator.

For the randomized delay task, trials with non-1.2 s delay lengths were used to calculate CD_choice_ (fit trials), always using the last 600 ms of the delay epoch as the time interval for calculation^65^. Population activity from trials with 1.2 s delays (test trials) were then projected along the CD_choice_ (**Fig. 3h**, *top left* and **Fig. 3i***, middle*). For visualization purposes, trials with all delay lengths (including fit trials) were projected along CD_choice_ in **Fig. 3i**, *top*.

### Reliability of choice decoding from CD_choice_

An ROC-AUC analysis was performed to estimate the reliability of choice decoding from projections along CD_choice_ on a single session basis (**EDFig. 2c**). For each session, equal numbers of correct left and rick lick trials were split into a training (70%) and testing set (30%). CD_choice_ was calculated from the activity of the neural population using training trials as described above. Activity from testing trials was projected along CD_choice_ and provided as input to a logistic regression model (fitglm() in MATLAB with distribution=’Binomial’, link=’logit’). The model was optimized to predict the animal’s choice on a particular trial from the delay epoch CD_choice_ activity. The model output was then passed into MATLAB’s perfcurve() function to obtain a receiver operating curve (ROC). Reliability of decoding was measured as the area under the ROC (AUC), shown in the inset of **EDFig. 2c**.

### Choice/context decoding from neural population or kinematic features

We trained logistic regression models to predict either animal choice (**Fig. 3b**) or behavioral context (**Fig. 4b**) from either kinematic features or single trial neural activity. The kinematic regressors were made up of the x and y positions and velocities of DeepLabCut-tracked features (tongue, jaw, and nose), as well as the tongue length, angle, and motion energy. The neural regressors were the firing rates of the units simultaneously recorded within each session. Equal numbers of correct left and right lick trials were used for this analysis. A separate model was trained at each time bin for both neural and kinematic decoding. Models were trained with ridge regularization and 4-fold cross-validation with 30% of trials held out for testing. Chance accuracy was obtained by shuffling choice/context labels across trials. Accuracy of the prediction was defined as:

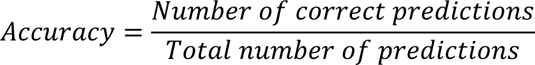

### Emergence of CD_choice_ selectivity/motion energy in randomized vs. fixed delay tasks

To find selectivity within the CD_choice_ projection (**Fig. 3h**, *top left*) we found the difference between the trial-averaged projection on right and left trials for each session. We then found the maximum selectivity value prior to the go cue and identified the first time point, relative to delay onset, for the selectivity trace to exceed 90% of the maximum value (**Fig. 3h**, *top right*). The same metric was found using session-averaged motion energy in **Fig. 3h**, *bottom panels*.

### Predicting projections along coding directions from kinematic features

We trained support vector machines (SVMs) to predict projections along CD_choice_, CD_ramp_, and CD_context_. Correct DR trials were used for predicting CD_choice_ and CD_ramp_ projections; correct DR and WC trials were used for predicting CD_context_. For predicting CD_context_, the regressors were made up of the x and y positions and velocities of DeepLabCut-tracked features (multiple points on the tongue, jaw, and nose taken from two camera angles), as well as the tongue length, angle, and motion energy (totaling 60 regressors). Only the sample and delay epoch were considered when predicting CD_choice_ and CD_ramp_ projections; because of this, any tongue-related metrics were not included as regressors in these analyses (totaling 55 regressors). Projections and kinematic features were first mean-centered and scaled to unit variance before input to the regression model. The previous B time bins of kinematic data were used to predict the current bin’s neural data (B=6, each bin is 5 ms). Models were trained with ridge regularization and 4-fold cross-validation. The models were tested on held-out testing data (40% of trials for CD_choice_ and CD_ramp_; tested on 30% of trials for CD_context_). To assess the prediction quality between projections and model predictions, we calculated the variance explained as:

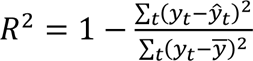

where *y_i_* is the value of the projection at time *t*, *ŷ_t_* is the prediction, and ȳ is the mean of the projection across all time.

When predicting projections along CD_choice_ during the randomized delay task, unregularized models were used (**Fig. 3h**). This was due to the small numbers of trials for each delay length which precluded the partitioning of data into 3 separate sets (training, validation, and testing trials). Instead, models were trained with linear regression and 4-fold cross-validation and tested on held-out data from each cross-validation fold. For assessing prediction quality between projections and model predictions, trials with 1.2 s delay epochs were used.

To understand which groups of kinematic features were most informative for predicting projections along CD_choice_, or CD_context_ (**EDFig. 3**), the absolute value of the beta coefficients for each kinematic regressor in the trained model was first obtained. For each feature group (jaw, nose, motion energy), the average beta coefficient across all regressors belonging to this kinematic feature group (8*B for jaw, 6*B for nose, and 1*B for motion energy; for example, the x and y positions and velocities for one point on the jaw on each camera totals 8 jaw regressors for each time bin, B) was calculated. The average beta coefficient for each feature group was then expressed as a fraction of the total of all beta coefficients.

### Subspace identification (trial-averaged data)

Delay and Response subspaces were identified following the general approach described in ref^40^. For identifying subspaces using trial-averaged data (**Fig. 5a**), neural activity was first trial-averaged separately for correct left and right trials. Activity across trial types was then concatenated, to produce a matrix ***X*** ∈ ℝ^(B*C)^ ^×^ ^N^, where B is the number of time bins, C the number of trial types, and N the number of units. Trial-averaged neural activity was soft-normalized (normalization factor=5) as described in ref^40^ and subsequently baseline subtracted (baseline: -2.4 to -2.2 s before go cue, during the ITI) The matrix, ***X***, was further divided into two matrices, ***X****_delay_* and ***X****_response_*. ***X****_delay_* contained activity from [-1,0] seconds relative to the go cue, and **X**_response_ contained activity from [0,1] seconds relative to the go cue. Delay and response subspaces were then identified by solving the following optimization problem:

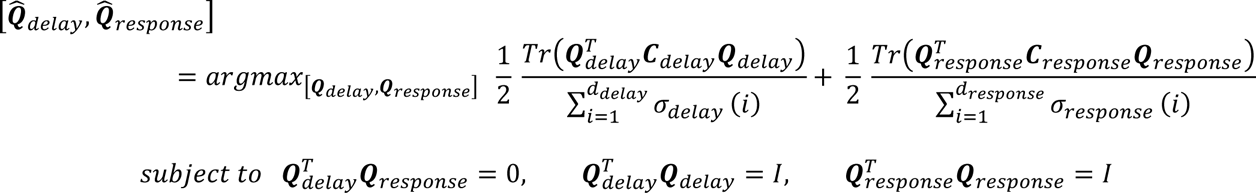

where ***C****_delay_* and ***C****_response_* are the covariances of ***X****_delay_* and ***X****_response_*, σ_*delay*_ and σ_*response*_ are the singular values of ***C****_delay_* and ***C****_response_*, and *d_delay_* and *d_response_* are the dimensionality of the subspaces. This optimization problem jointly identifies the subspaces that maximally contain the variance in neural activity during the delay and response epochs. The dimensionality of each space was chosen to be half the number of simultaneously recorded neurons, or twenty, whichever was smaller. Therefore, the dimensionality of the full population was equal to the dimensionality of the delay or response subspaces for sessions containing forty or fewer neurons. Optimization was performed using the manopt^107^ toolbox for MATLAB.

### Subspace identification (single-trial data)

For identifying subspaces using single-trial data (**Fig 5h**), single-trial neural activity was first binned in 5 ms intervals and smoothed with a causal Gaussian kernel with a half width of 35 ms. Each trial’s activity was subtracted by the average baseline activity across trials and scaled by the standard deviation across trials (baseline: -2.4 to -2.2 s before go cue, during the ITI). **X**_moving_ ∈ ℝ^B_p^ ^×^ ^N^ and **X** _stationary_ ∈ ℝ^B_n^ ^×^ ^N^ were defined using the single trial neural activity when the animal was moving or stationary (see ***Motion Energy***), respectively. B_n was the number of time bins during which the animal was annotated as stationary and B_p the number of time points annotated as moving. All correct and error trials from DR and WC contexts were used unless specified otherwise. Once the data was partitioned into moving and stationary time points, subspace identification was carried as described for trial-averaged data in the preceding paragraph:

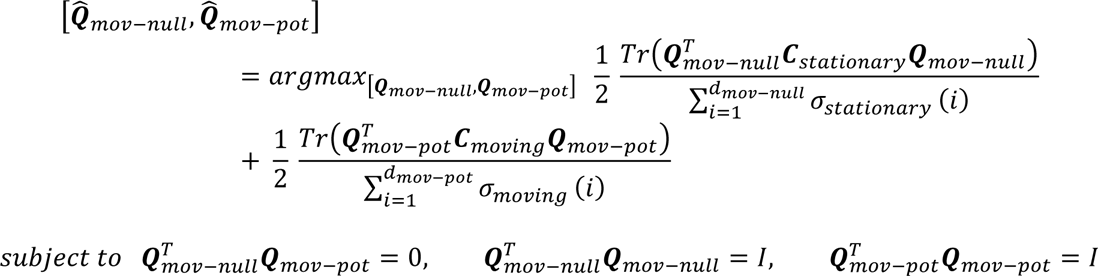

We quantified the normalized variance explained in a subspace as in refs^40,108^ (**Fig. 5d,k**):

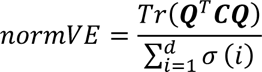

where **Q** is the subspace, **C** is the covariance of neural activity, and σ are the singular values of **C**. This normalization provides the maximum variance that can be captured by *d* dimensions.

Unit activity was reconstructed from movement-null and movement-potent subspaces according to the following equation:

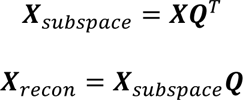

where ***X****_recon_* is the reconstructed neural activity, ***X****_subspace_* is the projected neural activity within the movement-null or movement-potent subspace, ***X*** is either single-trial or trial-averaged firing rates and ***Q*** is either the movement-null or movement-potent subspace.

Bootstrapped distributions of coding directions within the movement-null and movement-potent subspaces were obtained through two separate methods. For **Fig. 7** and **EDFig. 7a,c**, we first estimated movement-null and movement-potent subspaces per session and reconstructed neural activity from each subspace. Then, we performed the hierarchical bootstrapping procedure as described in ***Hierarchical bootstrapping***. For each iteration, we used the original neural activity to estimate the coding directions and then projected the reconstructed neural activity onto the coding directions. For **EDFig. 5b**, for each bootstrap iteration, the reconstructed neural activity itself was used to estimate the coding directions. Activity within each subspace was then projected along the respective coding directions. For **Fig. 7b,f**, coding directions were directly identified from the reconstructed neural activity, **X**_recon_, for the individual example sessions.

Subspace alignment for an individual unit was calculated as:

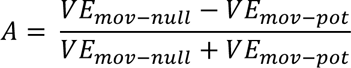

where VE is the variance explained of each individual unit by the movement-null or movement-potent subspace, or by the activity along a coding direction within the movement-null or movement-potent subspace. VE for a single unit was calculated as:

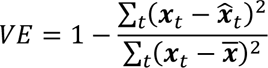

where ***x****_t_* is the trial-averaged firing rate, ***x^*** is the reconstructed trial-averaged firing rate, and *t* is the time bin.

To ask if the number of single units observed to be aligned to either the movement-null or movement-potent subspace was different than expected by chance, we compared the distributions to those obtained from randomly sampled subspaces as described in ref ^40^. Each element comprising a subspace was randomly sampled from a normal distribution with zero mean and unit variance but was biased by the covariance structure of the neural activity for a given session. Biasing the randomly sampled subspaces by the covariances controls for the unbalanced variance between stationary and moving time points (or between delay and response epochs). That is to say that the shuffled distributions take into account the relative amount of movement tuning across the neural population.

We sampled random subspaces, ***v_mov-null_*** and ***v_mov_*_-*pot*_** as follows:

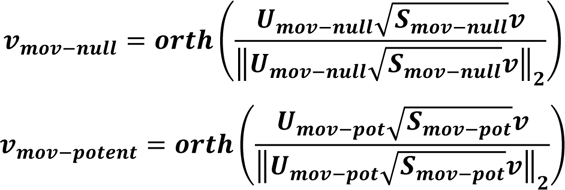

Where **U** and **S** are the left and right singular vectors of their associated covariance matrices, ***C_stationary_*** and ***C_moving_***, and ***v*** is a matrix whose elements are independently drawn from a normal distribution with zero mean and unit variance. As described above, covariance matrices ***C_stationary_*** and ***C_moving_*** are defined by neural activity during stationary and movement time points for single trial data. ***orth***(**A**) computes the orthonormal basis of a matrix **A**. Neural activity from each session was projected along the randomly sampled subspaces and alignment indices were calculated. This procedure was repeated 1000 times to generate null distributions, which represents the alignment indices of our data with random subspaces (**EDFig. 5)**. To generate *p*-values associated with **Fig. 5n**, **Fig. 7d,h**, **and Fig. 8d**, on each iteration, we computed the proportion of strongly tuned units (alignment ≥ 0.8 and alignment ≤ -0.8). This provided two distributions, one for alignment ≥ 0.8 and another for alignment ≤ -0.8. These chance distributions were separately fit with gaussian distributions using the fitgmdist() function in MATLAB. Then, *p*-values were computed as the probability of observing the proportion of tuned units we observe in the data from the fitted distributions.

Subspaces were identified using data from DR and WC correct and error trials (**Fig. 5,7**, and **8**). Control analyses were performed using separate sets of trials to assess if the movement-potent subspace erroneously contained movement-null-subspace dynamics. First, we estimated subspaces using WC trials only (**EDFigs. 4** and **8**). Second, we estimated subspaces using DR and WC trials, but restricted time points used to those in the response epoch only (**EDFigs. 4** and **8**). Both controls allowed us to estimate movement-null and movement-potent subspaces in the presence of uninstructed movements, but in the absence of planning dynamics. These controls thus allowed us to measure the degree to which the movement-potent subspace erroneously captured movement-null dynamics.

In a further attempt to validate that the movement-potent subspace is not inadvertently capturing movement-correlated internal dynamics, we identified movement-null and movement-potent subspaces using a two-stage PCA approach (**EDFig. 4**). The movement-null subspace was first identified as the dominant principal components (first 5 PCs) of the single-trial firing rates when mice are not moving. Second, the activity within the movement-null subspace was removed from single-trial firing rates. Then, the movement-potent subspace was calculated as the first 5 PCs of the single-trial residuals:

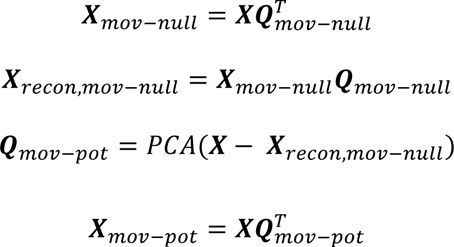

***X*** ∈ ℝ^(B*K)^ ^×^ ^N^ is the single-trial firing rates where B is the number of time points, K the number of trials, and N the number of units. ***Q***_mov-null_ is the first 5 PCs of the single-trial firing rates when animals are stationary, and ***X***_mov-null_ is the projection along those PCs. ***X***_recon,mov-null_ is the reconstructed neural activity obtained from the multiplication of activity within the movement-null subspace, ***X***_mov-null_, and the movement-null subspace, ***Q***_mov-null_. *PCA*(**Z**) indicates computing the PCs of the matrix **Z**. ***Q***_mov-pot_ is the first 5 PCs of the single-trial residuals obtained from subtracting ***X***_mov-null_ from the single-trial firing rates, ***X***. Finally, ***X***_mov-pot_ is obtained from projecting single-trial firing rates onto ***Q***_mov-pot_. ***X***_recon,mov-null_ contains activity that is explainable by the first 5 PCs in the absence of movement and, therefore, the residuals contain movement-related neural dynamics. Thus, performing PCA on these residuals provides a movement-potent subspace that is orthogonal to the movement-null subspace. This approach ensures dynamics observed during stationarity are contained within the movement-null subspace prior to assigning any dynamics to the movement-potent subspace. Therefore, it is more conservative in avoiding the mis-assignment of movement-correlated internal dynamics to the movement-potent subspace.

### CD_context_ selectivity on ‘High move’ and ‘Low move’ trials

For each session, DR and WC trials were first separated into ‘High move’ and ‘Low move’ trials. ‘High move’ trials were those where the average motion energy in the ITI was greater than the movement threshold for that session (see ***Motion energy***). ‘Low move’ trials were the remaining trials. The last 40 trials in each session were excluded from analysis to account for the decrease in uninstructed movements that are observed towards the end of behavioral sessions as animals become sated and disengaged. CD_context_ was calculated from the full population or from neural activity reconstructed from either subspace. Population activity from ‘High-move’ and ‘Low-move’ trials was then projected along CD_context_. Selectivity was defined as the trial-averaged projection along CD_context_ on DR trials minus the projection along CD_context_ on WC trials (**Fig. 8b,c**).

### Hierarchical bootstrapping

Projections along coding directions were obtained via a hierarchical bootstrapping procedure^109,110^ (**Figs. 2, 7** and **EDFigs. 7, 8** and **10)**. Pseudopopulations were constructed by randomly sampling with replacement M mice, 2 sessions per sampled mouse, 50 correct trials of each type, 20 error trials of each type, and 20 neurons. M is the number of mice in the original cohort. Bootstrapping was repeated for 1000 iterations. In each iteration, data derived from some individual mice (and sessions, trials, and neurons) will be overrepresented and some will be omitted. Average effects driven by small subsets of animals, sessions, trials, and/or units will be accompanied by large confidence intervals. For all results obtained through this bootstrapping procedure, mean and 95% confidence intervals (shaded area) of the bootstrap distribution are shown, except for selectivity (**Fig. 7a** and **EDFig. 7a**), where 5-95% confidence intervals are shown to indicate where projections significantly differ from zero (*p* < 0.05, one-sided test, bootstrap).

### Statistics

No statistical methods were used to determine sample sizes. All *t*-tests were two-sided unless stated otherwise.

## DATA AVAILABILITY

Primary and derived data described in this study will be made available on Figshare upon publication.

## CODE AVAILABILITY

MATLAB code for subspace identification is available at https://github.com/economolab/subspaceID. Custom MATLAB code used for analyses will be made available on Github upon publication.

## AUTHOR CONTRIBUTIONS

MAH, JEB, and MNE conceived of the project. MNE and CC supervised research. MAH, JEB, and MNE designed experiments. MAH, JEB, JLUN, and EKH performed experiments. MAH, JEB, and MNE analyzed data. MAH, JEB and MNE wrote the manuscript.

## ACKNOWLEDGMENTS

The authors thank Karel Svoboda, Brian DePasquale, Shaul Druckmann, and Ben Scott for helpful discussions. We thank Tim Wang for helpful comments on the manuscript. We thank John Jiang for help with cranial window surgeries. This work was supported by the Whitehall Foundation, the Klingenstein Fund, the Simons Foundation, and NIH R01NS121409.

## Extended Data

**Extended Data Fig. 1.**
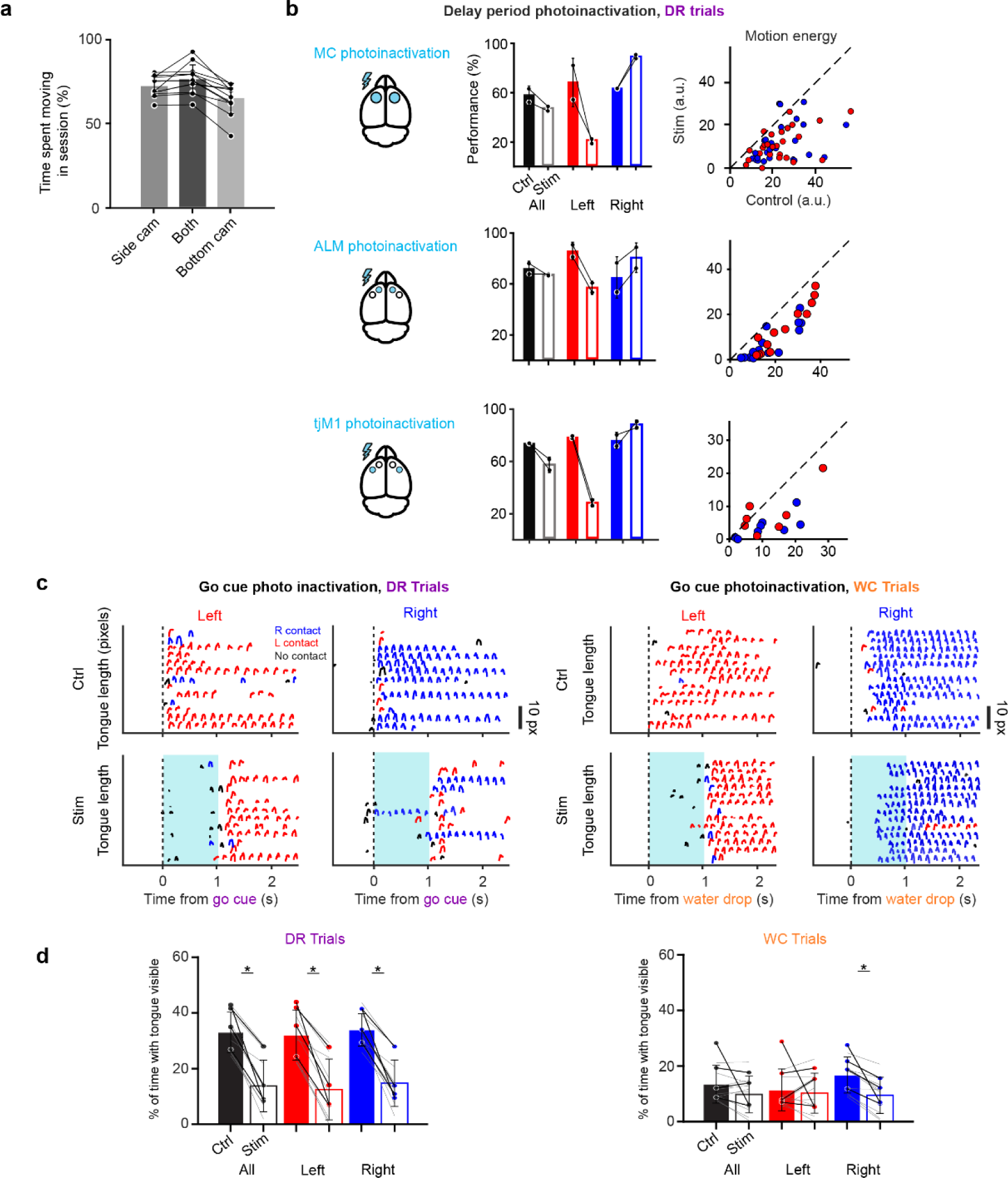
Region-specific photoinactivation of ALM and tjM1. **a.** Percentage of time spent moving (see **Methods**) in a session determined by video recordings using the side view, bottom view, and both views. Points indicate each of the 12 randomly selected sessions used for this analysis. **b.** Effect of delay epoch photoinactivation on behavioral performance (*middle column*) and uninstructed movements (*right column*) when photoinactivation was directed to the MC (ALM + tjM1; *top row*; same data as Fig. 1g**-i**, *n* = 14 sessions, 2 mice), the ALM (*middle row*, *n* = 15 sessions, 2 mice), and the tjM1 (*bottom*, *n* = 9 sessions, 2 mice). Photoinactivation of ALM and tjM1 led to similar behavioral impairment and reduction in uninstructed movements, with larger effects observed with MC (ALM + tjM1) photoinactivation. **c.** Tongue length during control and go cue photoinactivation trials for delayed-response (*left*) and water-cued (*right*) contexts. Blue traces indicate right lickport contacts, red traces indicate left contacts, and black traces indicate no contact. Vertical dashed line indicates go cue or water drop onset. Blue shaded region indicates photoinactivation period. **d.** Percentage of time with tongue visible during photoinactivation period for DR trials (*left*) and WC trials (*right*). Each colored point indicates mean value for an animal (*n* = 4 animals), individual animals are connected by black lines. Light gray lines denote individual sessions (*n* = 10 sessions). Bars are the mean across all sessions. Asterisks denote significant differences (*p* < 0.05) between control and photoinactivation trials (Percent reduction on all DR trials: 19 ± 7%, mean ± s.d., *p* = 1.6e-05; DR left trials: 20 ± 8%, *p* = 3.0e-05; DR right trials: 20 ± 7%, *p* = 1.6e-05; All WC trials: 4 ± 8%, p=0.154; WC left trials: 1% ± 9%, p=0.702; WC right trials: 7% ± 5%, p=0.002; paired *t-*test, *n*=10 sessions). Error bars indicate standard deviation across sessions. In WC trials, tongue protrusion was only significantly impaired on one trial type, while ability to successfully contact the lickport was impaired in all conditions (see Fig. 1c).

**Extended Data Fig. 2.**
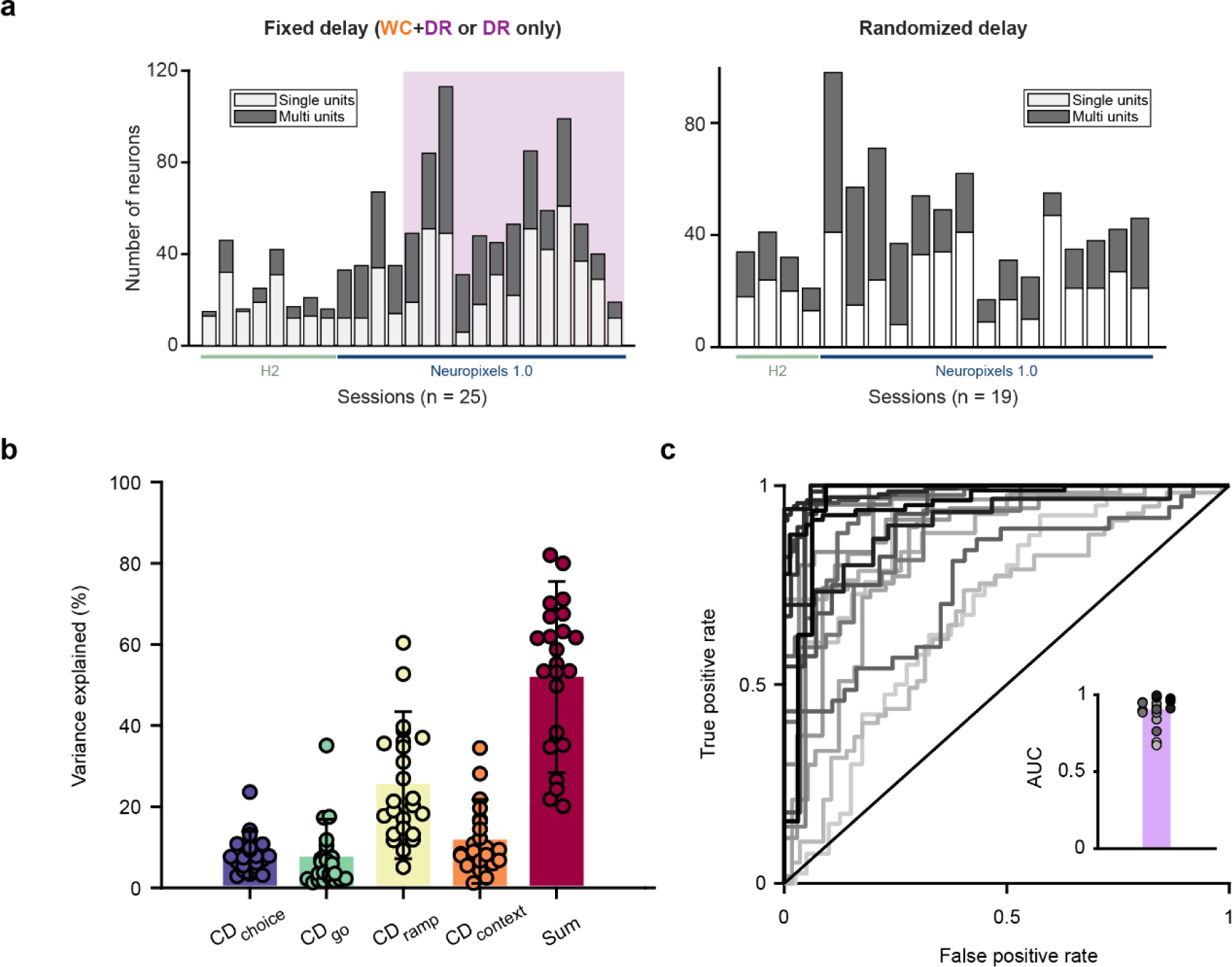
Session-by-session statistics. **a.** Number of recorded single- and multi-units per session for the fixed delay task (*left*) or the randomized delay task (*right*). *Left*, Purple shaded region indicates sessions in which animals only performed presented with DR trials. Green and blue bars underneath plots indicate the probe type used for a given session **b.** Variance explained of trial-averaged neural activity by each coding direction. The coding directions were calculated using neural activity from individual sessions (n=25). Bar height represents the mean across sessions and error bars indicate standard deviation across sessions. **c.** Receiver operating curves (ROC) demonstrating choice decoding accuracy from delay epoch CD_choice_ projections across all individual sessions (see **Methods**). Inset: area under the ROC curve (AUC). Bar height represents the mean across sessions and points indicate sessions.

**Extended Data Fig. 3.**
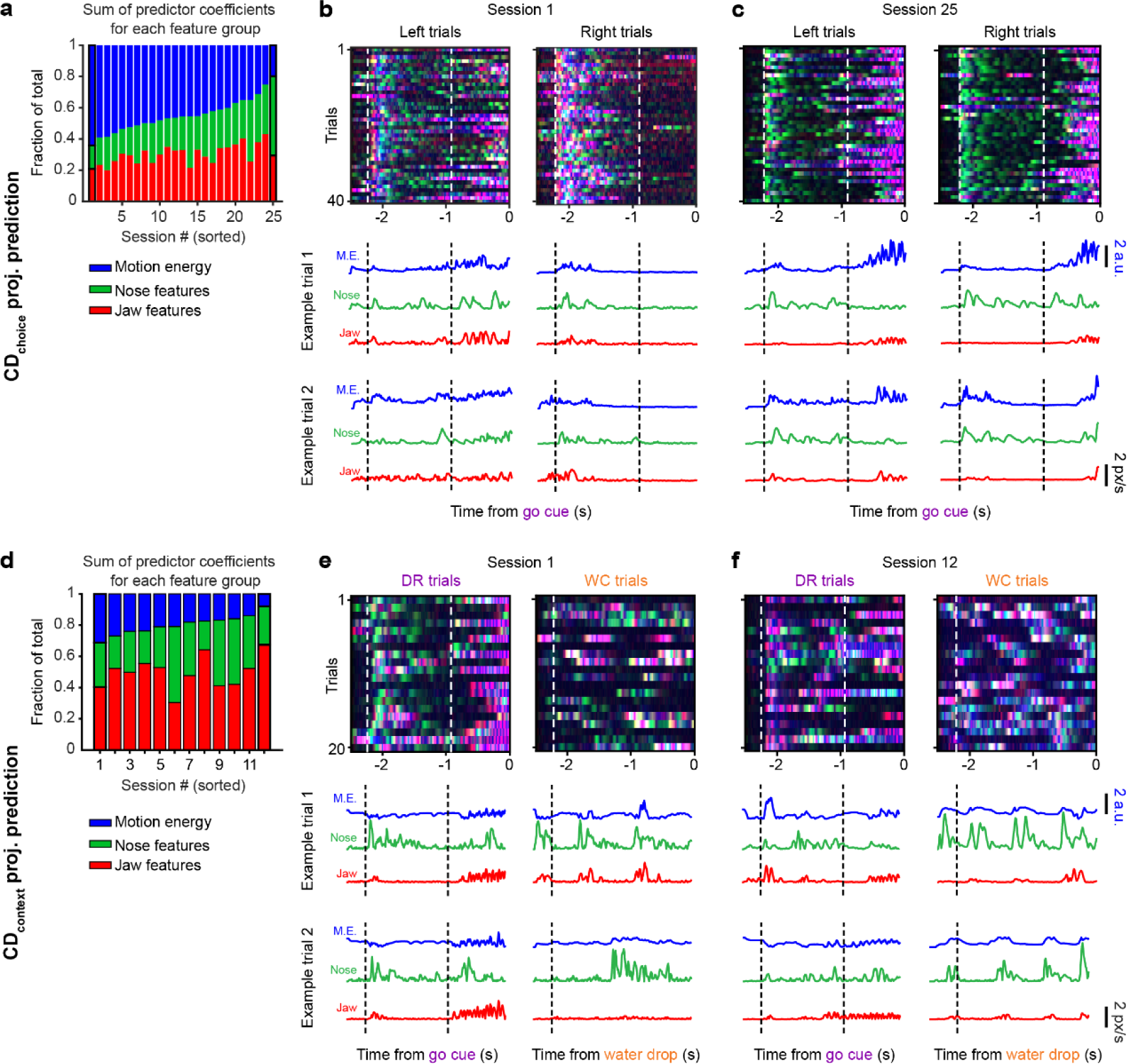
Session-by-session variability in the relationship between kinematic features and putative cognitive dynamics. **a.** Regression weights for each group of kinematic predictors of CD_choice_ projections, as a fraction of all predictor coefficients (see **Methods**). Sessions are sorted in descending order by motion energy fraction. Outlined bars indicate example sessions shown in (b) and (c). **b.** Example session where motion energy made up the largest fraction of regression weights for predicting CD_choice_ projections. *Top*, overlayed jaw/nose speed or motion energy for a subset trials. *Bottom*, two example trials of kinematic feature trajectories. **c.** Same as (b) but for an example session where jaw and nose features made up a larger fraction of regression weights. **d.** Same as (a) but for predicting CD_context_ projections. **e, f.** Same as (b) and (c) but for two example sessions with different regression weight fractions for predicting CD_context_ projections.

**Extended Data Fig. 4.**
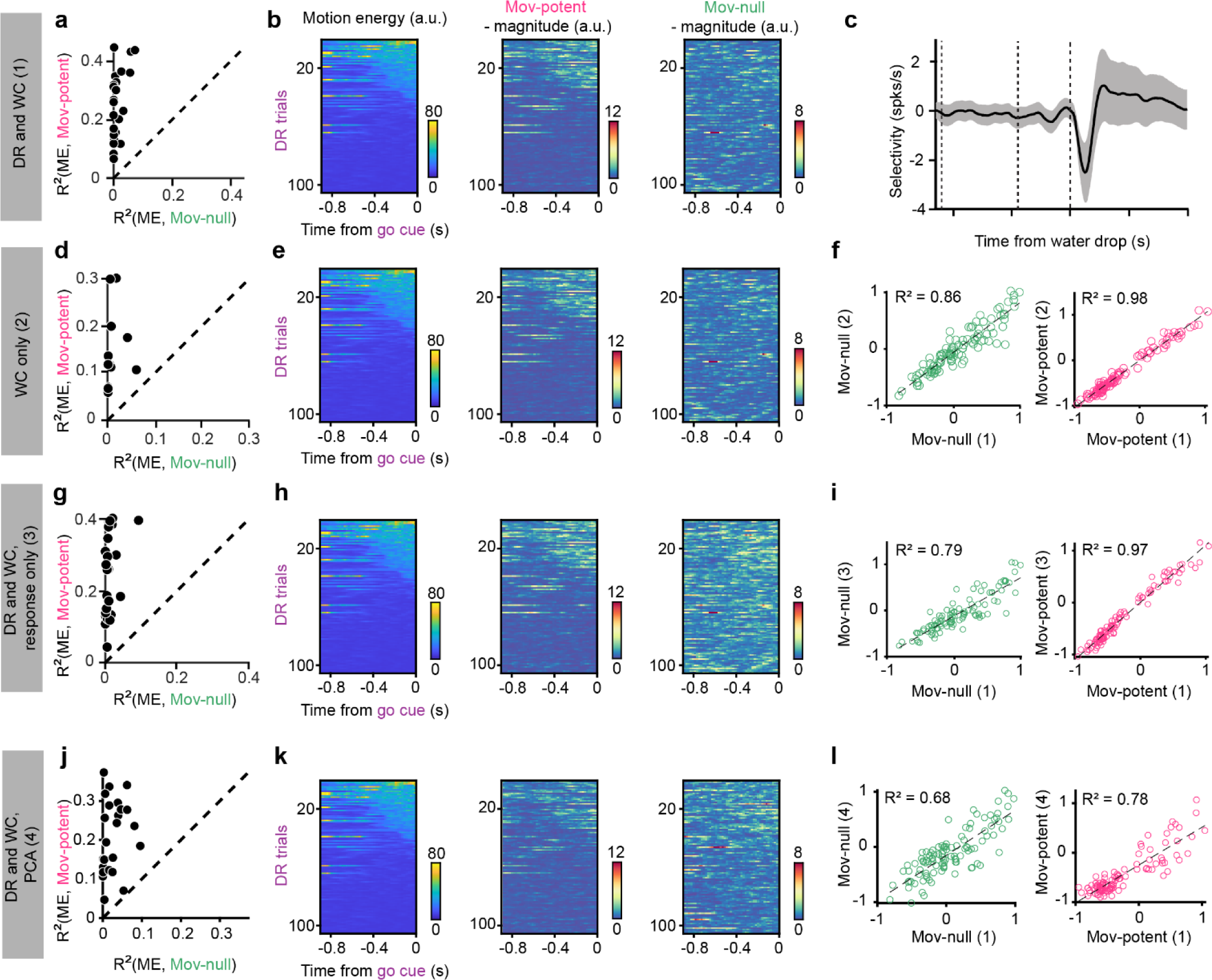
Control analyses for subspace decomposition. **a,b.** Movement-null and movement-potent subspaces estimated as in Fig. 5g**-l** using DR and WC trials. **a.** Variance explained (R^2^) of motion energy by the sum squared magnitude of activity in the movement-null and movement-potent subspaces on single trials. Each point is the mean across trials for a session. **b.** *Left*, motion energy on single trials for an example session. *Middle*, sum-squared magnitude of activity in the movement-potent subspace. *Right*, sum-squared magnitude of activity in the movement-null subspace. Trials sorted by average delay epoch motion energy. **c.** Selectivity (left vs. right) of the neural population during WC trials. Mean and 95% CI across sessions shown. **d,e.** Same as (a,b) but estimating movement-null and movement-potent subspaces using WC trials only. **f.** Normalized magnitude of activity in the movement-null subspace (*left*) movement-potent subspace (*right*) when estimated using DR and WC trials as in (a,b), versus when estimated using WC trials only as in (d,e). Circles are average activity per trial for an example session. **g,h.** Same as (a,b), but estimating movement-null and movement-potent subspaces using data restricted to the response epoch of DR and WC trials. **i.** Magnitude of activity in the movement-null subspace (*left*) or movement-potent subspace (*right*) when estimated using DR and WC trials as in (a,b) versus when estimated using data from only the response epoch of DR and WC trials as in (g,h). Circles are average activity per trial for an example session. **j,k.** Same as (a,b), but estimating the movement-null and movement-potent subspaces using a two-stage PCA approach (see **Methods**). This approach is conservative in avoiding the mis-assignment of cognitive dynamics that correlate in time with movement to the movement-potent subspace. **j.** Magnitude of activity in the movement-null subspace (*left*) or movement-potent subspace (*right*) when estimated using DR and WC trials as in (a,b) versus when estimated using the two-stage PCA approach as in (j,k). Circles are average activity per trial for an example session.

**Extended Data Fig. 5.**
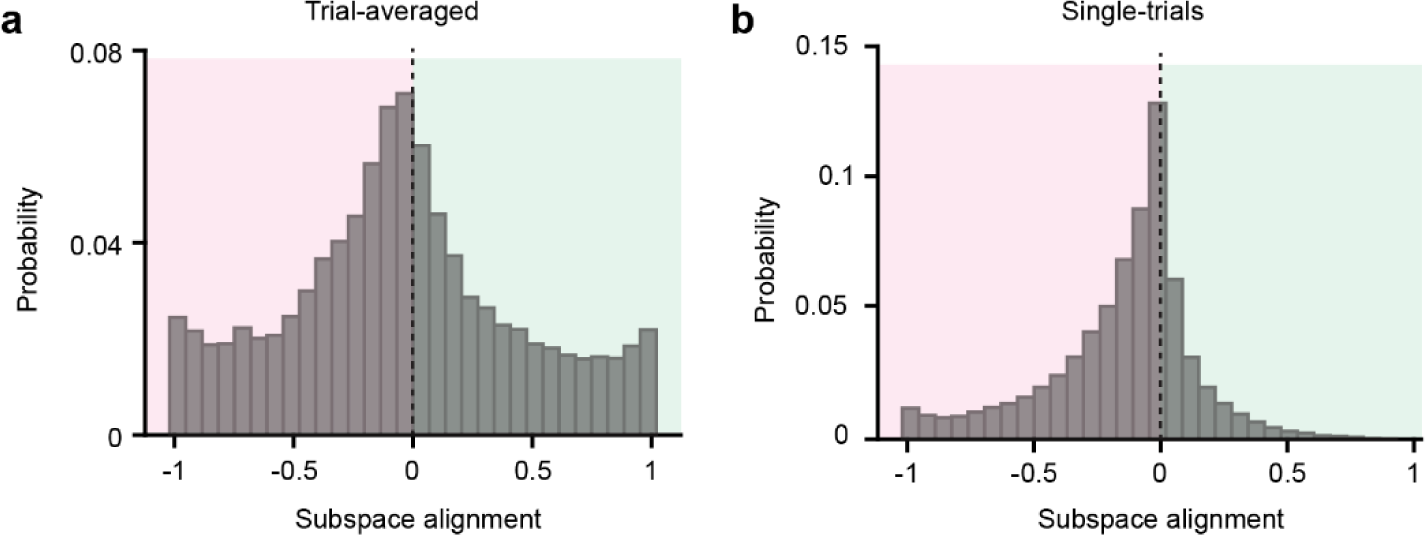
Alignment of single-units to random subspaces. Random subspaces were constructed by independently and identically drawing from a normal distribution with zero mean and unit variance. Each random subspace was then biased towards the covariance structure of the actual data (see **Methods**). **a.** Null distributions of alignment indices for trial-averaged data. **b.** Null distributions of alignment indices for single-trial data. Null alignment distributions are skewed towards the movement-potent subspace due to the unbalanced variance between delay and response epochs (a) or between stationary and movement time points (b), reflecting the strong movement tuning of many units.

**Extended Data Fig. 6.**
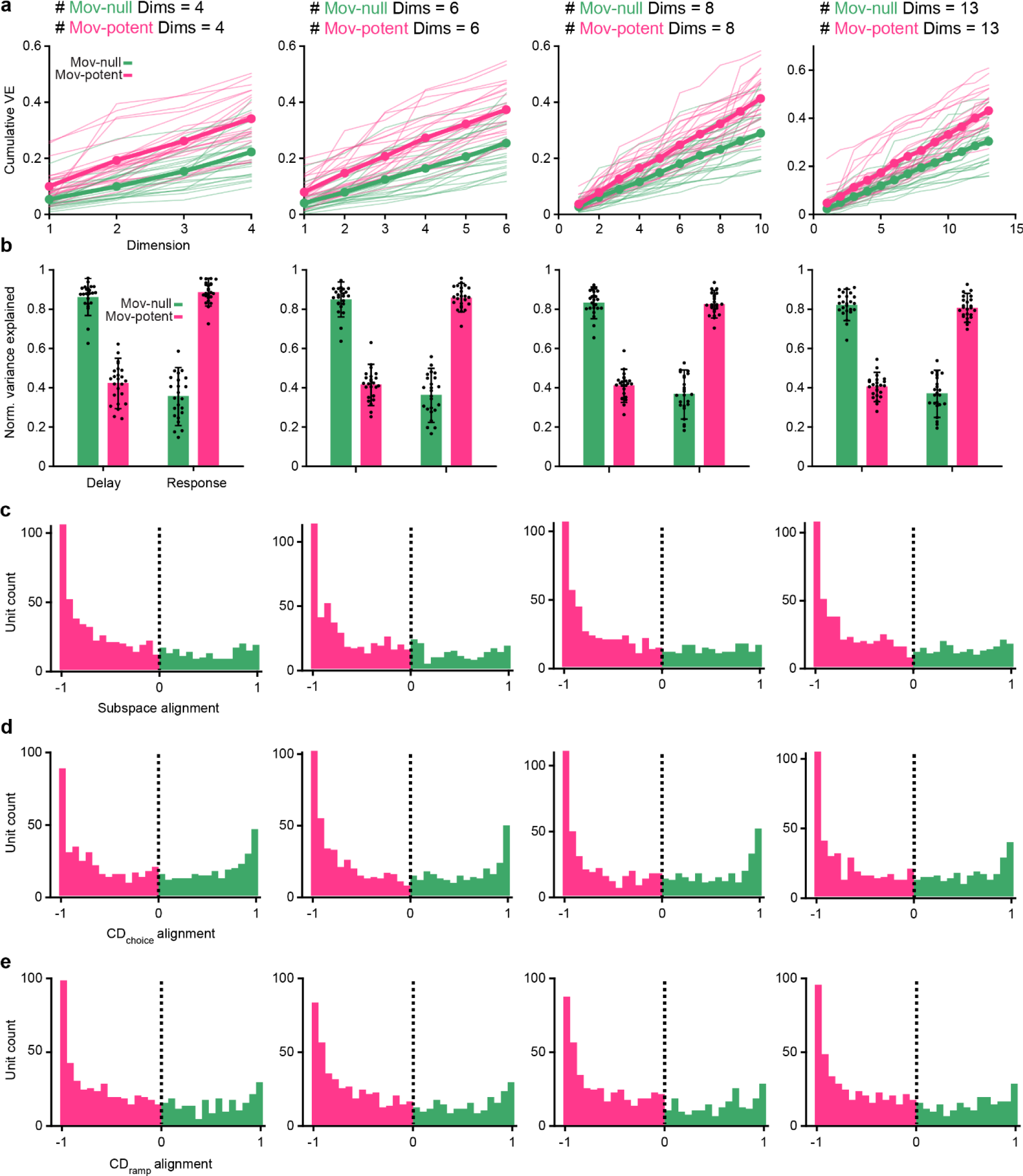
Varying dimensionality of subspaces. Analyses were repeated while varying the dimensionality of movement-null and movement-potent subspaces. Each subspace was constrained to be 4 (*left*), 6 (*middle left*), 8 (*middle right*), or 13 dimensions (*right*). **A.** Cumulative variance explained of the neural activity by the activity in movement-null and movement-potent subspaces. Bold lines and points indicate mean across sessions. Thin lines represent single sessions **b.** Normalized variance explained of neural activity during the delay or response epoch by the activity in movement-null and movement-potent subspaces. Points indicate sessions, bar height indicates mean across sessions, and error bars indicate standard deviation across sessions (n=25 sessions). **c-e.** Subspace (**c**), CD_choice_ (**d**), and CD_ramp_ (**e**) alignment distributions when varying dimensionalities of each subspace.

**Extended Data Fig. 7.**
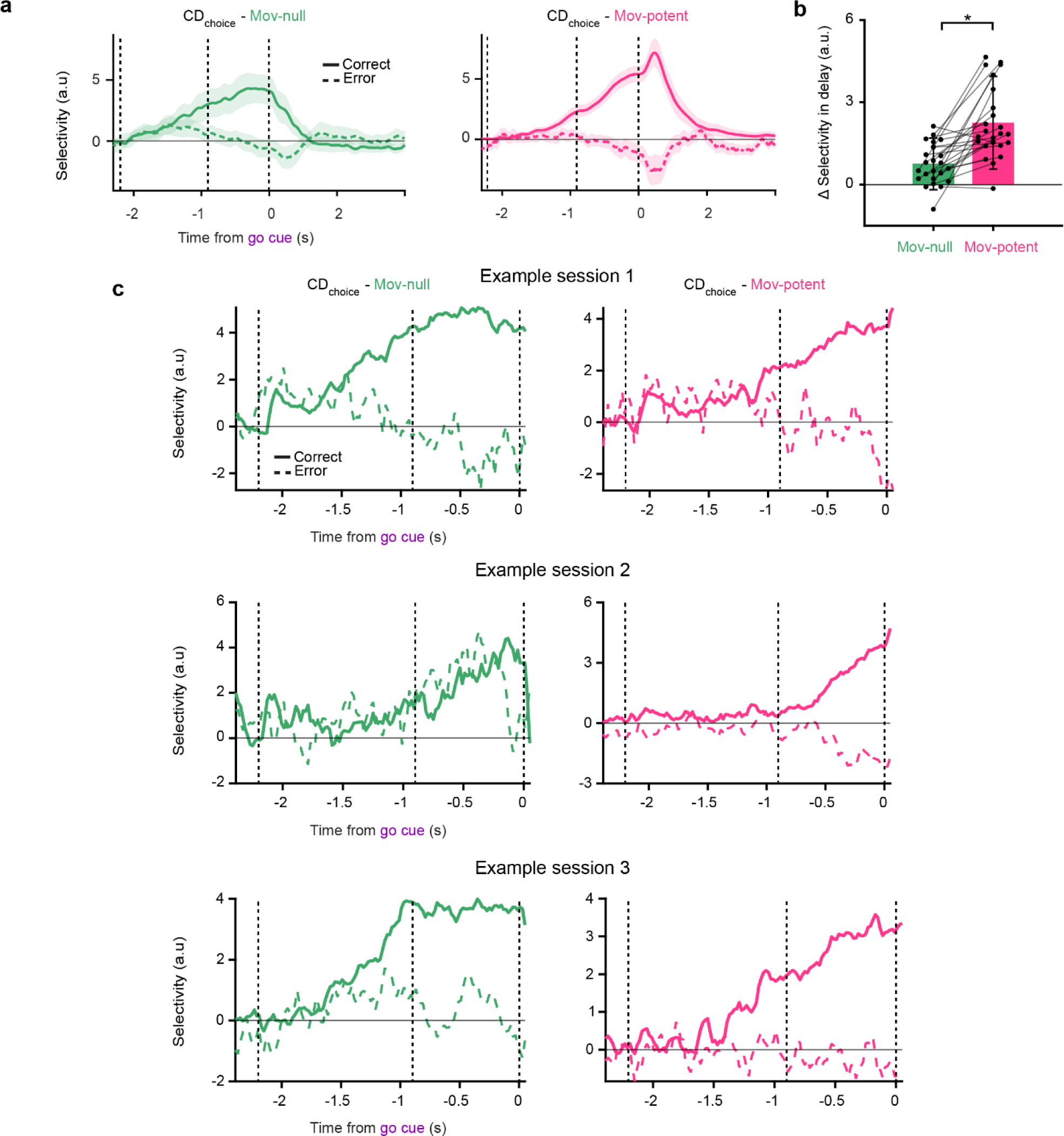
Projections along movement-null and movement-potent components of CD_choice_. **a.** Same data as in Fig. 7a except all time in trial shown to highlight activity during the response epoch. Selectivity (projections onto CD_choice_ on lick-right trials minus projections on lick-left trials) of movement-null (*left*) and movement-potent (*right*) subspace activity. Mean and 5-95% CI of the bootstrap distribution for correct (*solid*) and error (*dashed*) trials shown. **b**. Change in selectivity between the last 100ms of the delay epoch and the last 100ms of the sample epoch in movement-null and movement-potent components of projections along CD_choice_ (Movement-potent: 2.25 ± 1.57, mean ± s.d., Movement-null: 0.76 ± 0.8, *p*=1 x10^-5^, paired *t*-test, *n* = 25 sessions). Points indicate individual sessions, bar height indicates mean across sessions, and error bars indicate standard deviation across sessions. **c.** Three example sessions from three different mice depicting selectivity along CD_choice_ as in Fig. 7a. Solid lines denote the mean projection on correct trials and dashed lines denote the mean projection error trials.

**Extended Data Fig. 8.**
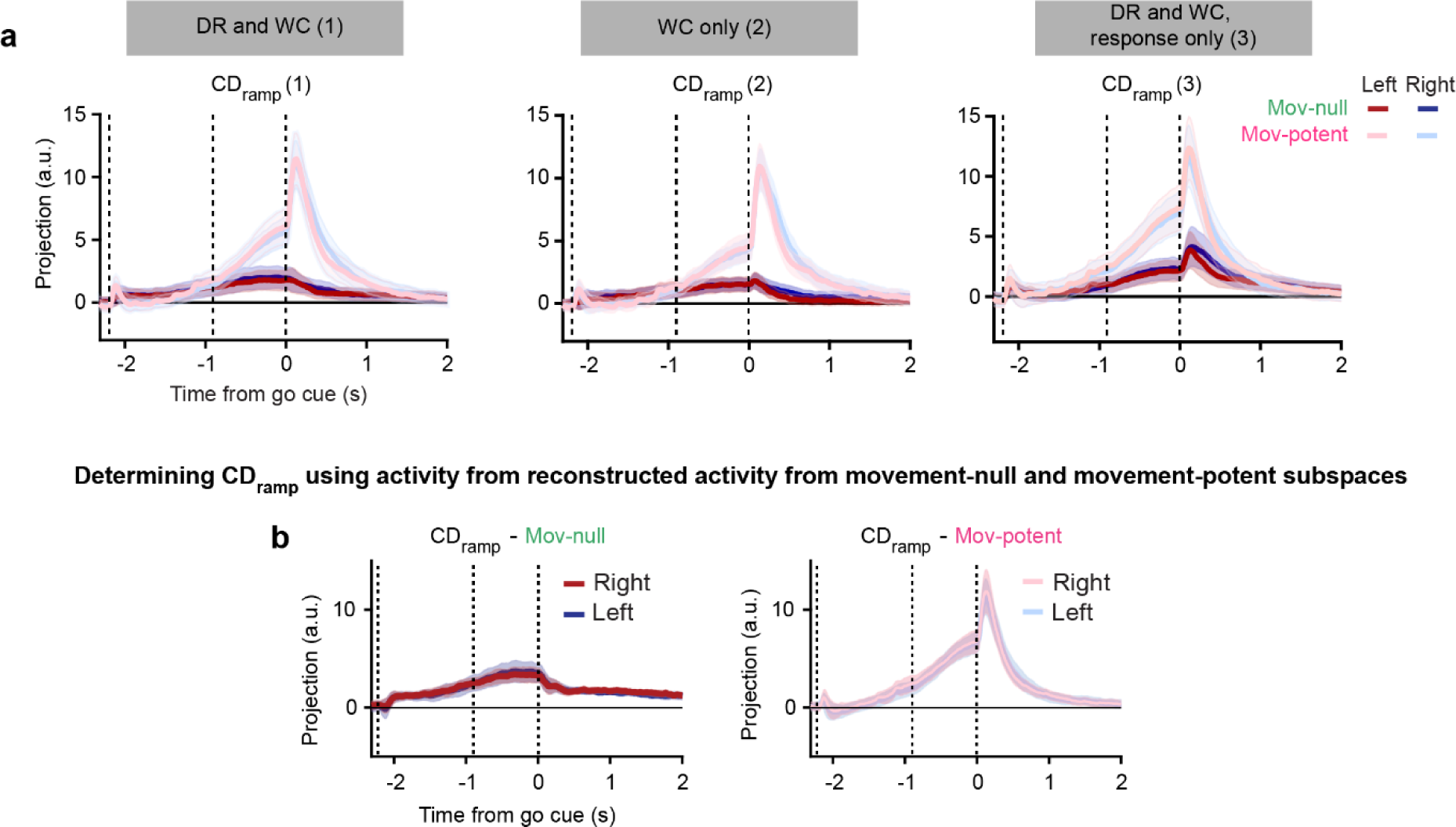
Within-subspace CD projections using variations on procedure to determine subspaces. **a.** Projections of movement-null and movement-potent subspace activity along CD_ramp_ for each of three analytical variations. Movement-null and movement-potent subspaces were identified using both DR and WC trials (*left*), WC trials only (*middle*), and the response epoch of DR and WC trials (*right*). Mean and 95% CI of bootstrap distribution shown.. **b.** Projections along movement-null (*left*) and movement-potent (*right*) components of CD_ramp_ when determined from activity within each subspace individually, rather than from the full neural population. Mean and 95% CI of bootstrap distribution shown.

**Extended Data Fig. 9.**
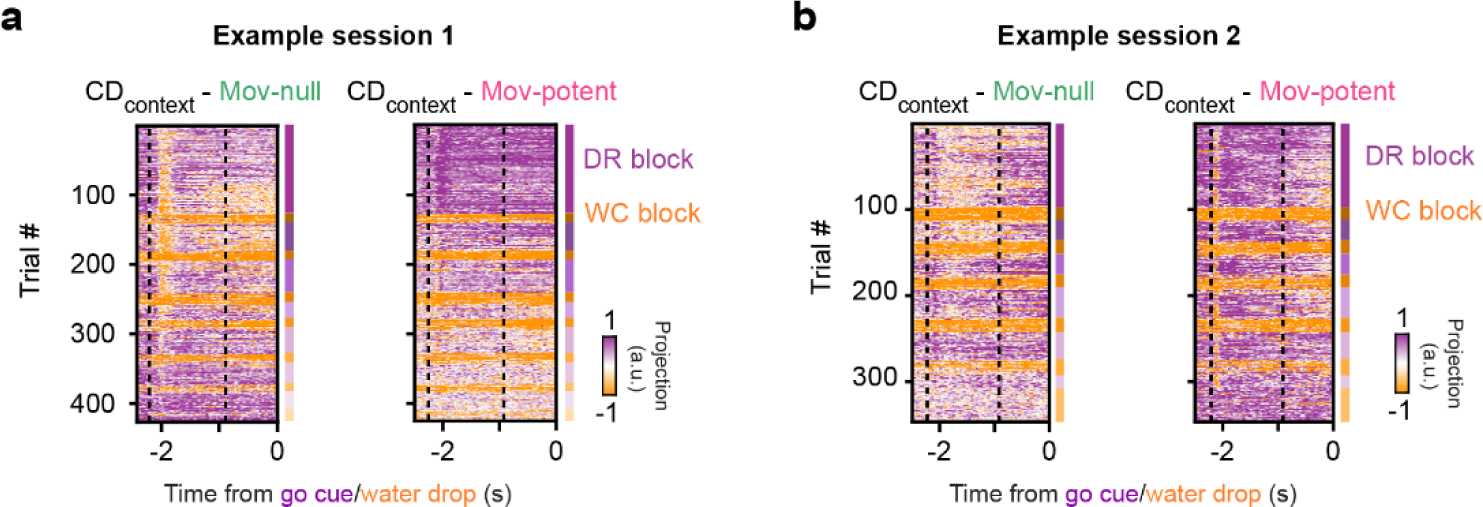
Encoding of context in both the null and potent subspaces tracks block-wise task structure. **a.** Heatmap of single-trial projections of null and potent subspace activity along CD_context_ for an example session. The chronological DR or WC block within the session is denoted by differently shaded purple and orange rectangles, respectively, on the right of each plot. **b.** Same as (a) but for another example session, from a different animal.

**Extended Data Fig. 10.**
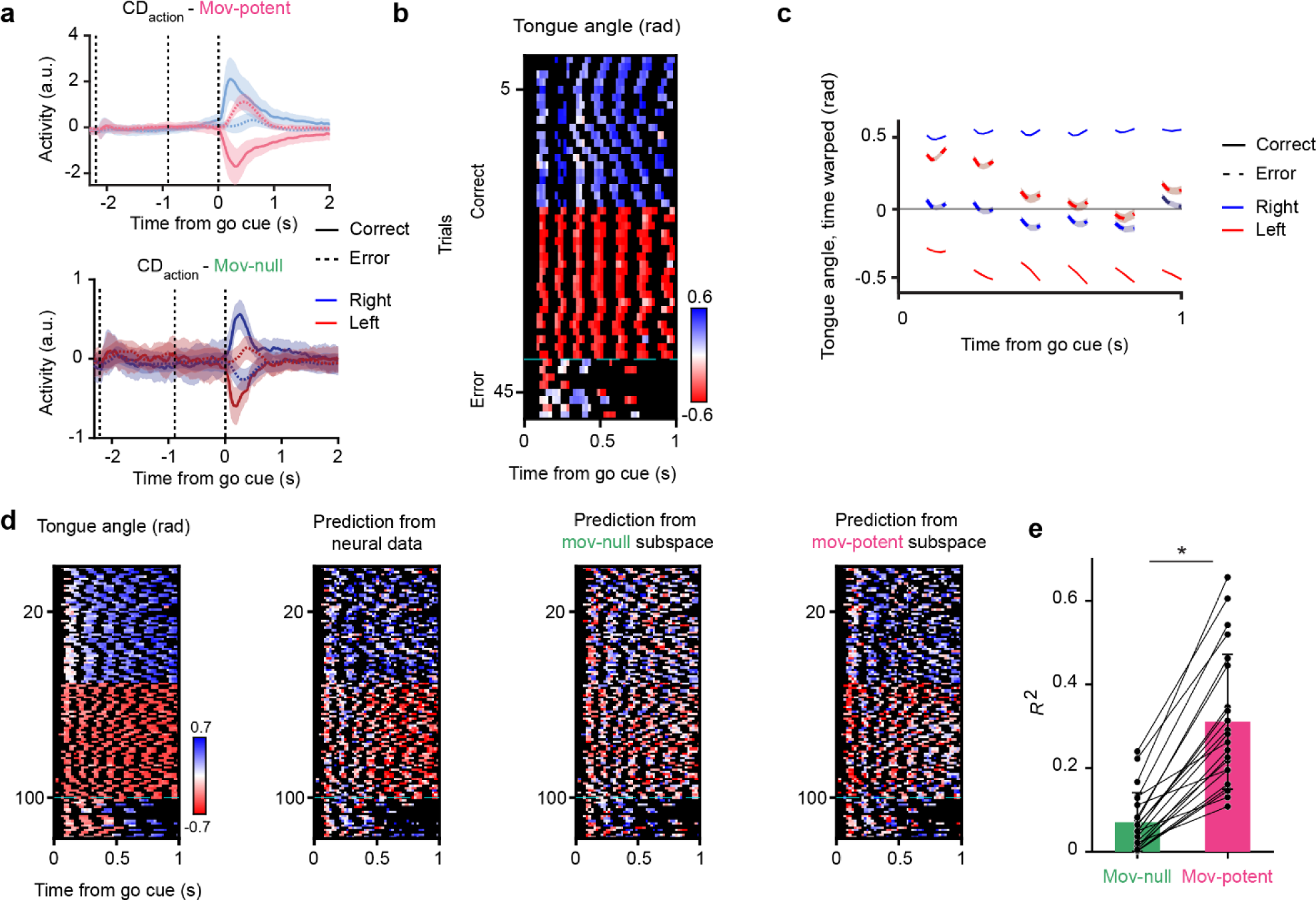
Relationship between tongue angle and neural activity in the movement-null and movement-potent subspaces. **a.** Projections along movement-potent (*top*) and movement-null (*bottom*) components of CD_action_. Correct trials shown in solid lines and error trials shown in dashed lines. **b.** Tongue angle for an example session for correct and error trials Black values indicate tongue not visible. **c.** Tongue angle on correct and error right and left trials. Tongue angle was linearly time warped to allow for averaging over trials and sessions. Mean and s.e.m. across sessions shown. **d.** Tongue angle (*left*) and predictions from the full population neural activity (*middle left*), null subspace activity (*middle right*), and potent subspace activity (*right*) for an example session. **e.** Variance explained (R^2^) of tongue angle by prediction from movement-null (*green*) and movement-potent (pink) subspaces. Asterisks denote significant differences between predictions from null and potent subspaces (*p*=2 x10^-8^, paired t-test, n=25 sessions).

**Supplementary Movie 1 – Uninstructed movements vary in their identity and timing.** Example trials in which uninstructed movements vary in their identity (across rows) and timing (across columns). Traces represent the y-position of the feature within the video frame. All example trials taken from the same mouse and session.

